# Size-dependent increase in RNA Polymerase II initiation rates mediates gene expression scaling with cell size

**DOI:** 10.1101/754788

**Authors:** Xi-Ming Sun, Anthony Bowman, Miles Priestman, Francois Bertaux, Amalia Martinez-Segura, Wenhao Tang, Dirk Dormann, Vahid Shahrezaei, Samuel Marguerat

**Affiliations:** MRC London Institute of Medical Sciences (LMS), Du Cane Road, London W12 0NN, UK; Institute of Clinical Sciences (ICS), Faculty of Medicine, Imperial College London, Du Cane Road, London W12 0NN, UK; Department of Mathematics, Faculty of Natural Sciences, Imperial College London, London SW7 2AZ, UK

## Abstract

Cell size varies during the cell cycle and in response to external stimuli. This requires the tight coordination, or “scaling”, of mRNA and protein quantities with the cell volume in order to maintain biomolecules concentrations and cell density. Evidence in cell populations and single cells indicates that scaling relies on the coordination of mRNA transcription rates with cell size. Here we use a combination of single-molecule fluorescence *in situ* hybridisation (smFISH), time-lapse microscopy and mathematical modelling in single fission yeast cells to uncover the precise molecular mechanisms that control transcription rates scaling with cell size. Linear scaling of mRNA quantities is apparent in single fission yeast cells during a normal cell cycle. Transcription rates of both constitutive and regulated genes scale with cell size without evidence for transcriptional bursting. Modelling and experimental data indicate that scaling relies on the coordination of RNAPII transcription initiation rates with cell size and that RNAPII is a limiting factor. We show using real-time quantitative imaging that size increase is accompanied by a rapid concentration independent recruitment of RNAPII onto chromatin. Finally, we find that in multinucleated cells, scaling is set at the level of single nuclei and not the entire cell, making the nucleus the transcriptional scaling unit. Integrating our observations in a mechanistic model of RNAPII mediated transcription, we propose that scaling of gene expression with cell size is the consequence of competition between genes for limiting RNAPII.

## INTRODUCTION

Gene expression is coordinated with cell size in order to maintain biomolecule concentrations. Understanding the mechanisms that mediate this coordination, called hereafter “scaling”, is a fundamental and intriguing problem in cell biology [1, 2]. Messenger RNAs (mRNA) and proteins are synthesised from the cell DNA genome, which is one of the few cellular components that do not scale with size. Because cell volume increases exponentially and mRNA half-lives are typically short, constant rates of mRNA or protein production cannot lead to gene expression scaling. Recent work, in yeast [3], animal [4, 5] and plant cells [6] has shown that mRNA synthesis rates instead are coordinated globally with cell size and are a major mechanism of scaling. Conversely, mRNA degradation seems to be mostly unconnected to scaling [3,4,6], although evidence suggests that degradation rates are adjusted early after budding yeast asymmetric division [7] and when growth rate changes [8, 9]. Scaling is pervasive and only few mRNAs are able to deviate from its regulation [10–12]. Interestingly, two of them have been found to participate in the control of size homeostasis [10, 11].

What could be the molecular mechanism behind transcription scaling with cell size? For a gene with an active promoter mRNA numbers follow a Poisson distribution [13]. Transcription is however often discontinuous and periods of RNA synthesis or ‘bursts” alternate with periods of promoter inactivity [14]. Work in single mammalian cells has shown that scaling of mRNA numbers results from a coordination of the size of the transcription bursts with cell volume and not from their frequency [4]. This is compatible with the increased RNAPII occupancy observed genome-wide in large fission yeast cell cycle mutants [3]. This also indicates that the mechanism behind scaling may not be related to activation of transcription but rather to the efficiency of an active promoter. Critically, transcription is a complex process and is regulated at many levels including, RNAPII initiation, pause/release, elongation, and termination [15–18]. Which of these processes is coordinated with cell size to mediate scaling remains unclear.

How could a complex set of molecular reactions such as transcription become more efficient as cell size increases? In an elegant experiment Padovan-Merhar and colleagues fused cells of small and large size and found that the number of mRNAs from a gene encoded in the small cell genome would increase in response to the large cell environment, however its concentration was almost halved. This suggests that scaling responds to both cell volume and DNA content [4]. This is consistent with the observation that the cell synthesis capacity is split between genome copies in diploid budding yeast cells [10]. Importantly, changes in gene numbers, in the case of increased ploidy for instance, are associated with overall cell size increase in many organisms [1]. This indicates that the number of genes present in a cell is linked to its volume and number of macromolecules. This also suggests that the cell’s overall synthetic capacity could be limiting and have a determining role in setting its size [19]. It is therefore likely that scaling depends on a limiting factor involved in transcription but its identity and regulation with the cell volume is not known [19, 20].

In this study, we measured gene expression in over 20,000 single cells of the fission yeast *Schizosaccharomyces pombe* by single molecule *in situ* hybridisation (smFISH, **Tables S1-3**) [21]. We combined these data with agent-based models of growing and dividing cells [22–24], stochastic models of gene expression [13,25,26] and Bayesian inference [27–33] to investigate the quantitative parameters of gene expression that mediate scaling. This integrative approach enabled us to determine which part of the transcription process is scaling with cell size and which molecular event connects transcription with the cell volume.

## RESULTS

### Gene expression scaling is a single cell attribute of constitutive and inducible gene expression

We first confirmed that scaling of gene expression with cell size was an attribute of single fission yeast cells during rapid proliferation. To do this, we measured mRNA levels of 7 constitutively expressed genes in wild-type haploids (*wt*), and in conditional mutants of the Wee1 (*wee1-50*) and Cdc25 (*cdc25-22*) cell cycle regulators by smFISH (**Figure 1A, Table S1**). At semi-permissive temperature, mutant cells divide at smaller and larger sizes than *wt* respectively (**Figure S1A**) [3]. Consistent with population microarray data, mean numbers of the 7 mRNAs measured in single cells scale with the average size of the three strains (**Figure 1B**) [3].

**Figure 1:**
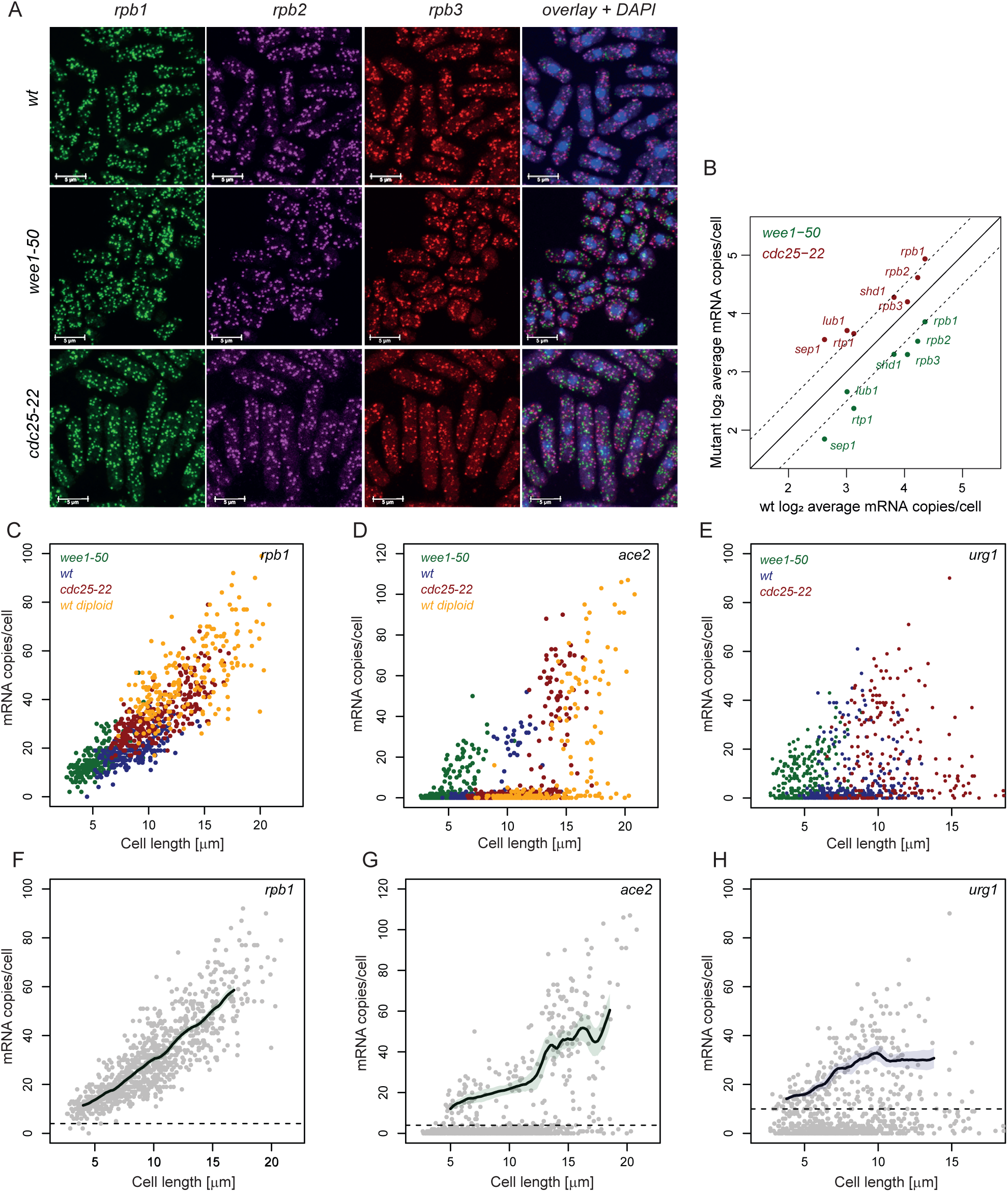
Gene expression scaling is a single cell attribute of constitutive and inducible gene expression. **A**. Representative images of an smFISH experiment. The *rpb1*, *rpb2*, and *rpb3* mRNA are labelled in *wee1-50*, *cdc25-22*, and *wt* cells. Overlay of the three channels and DAPI staining of DNA are shown in the last column. The white scale bar represents 5µm. **B**. Log2 average copies/cell of 7 mRNAs in wee1-50 (green) and cdc25-22 (red) cells plotted against log2 average copies/cell of the same mRNAs in *wt* cells. mRNA common names are shown on the figure. Plain line shows equality and dotted lines values 2 fold up or down. **C**. *rpb1* mRNA copies/cell plotted against cell length for *wee1-50* (green), *wt* (blue), *cdc25-22* (red) and *wt* diploid cells (orange). **D**. *ace2* mRNA copies/cell plotted against cell length for *wee1-50* (green), *wt* (blue), *cdc25-22* (red) and wt diploid cells (orange). **E**. *urg1* mRNA copies/cell plotted against cell length for *wee1-50* (green), *wt* (blue), *cdc25-22* (red) cells. Cells were analysed 3h after addition of uracil to the medium. **F**. same as C. with all the cells in grey. The solid line shows median counts in running windows sampled from 100 bootstrap samples of the experimental data and the shaded area represents 95% confidence intervals. Only cells above the dotted lines were considered as expressing the mRNA and were used for the running window analysis. **G**. same as D. with all the cells in grey. The solid line shows median counts in running windows sampled from 100 bootstrap samples of the experimental data and the shaded area represents 95% confidence intervals. Only cells above the dotted lines were considered as expressing the mRNA and were used for the running window analysis. **H**. same as E. with all the cells in grey. The solid line shows median counts in running windows sampled from 100 bootstrap samples of the experimental data and the shaded area represents 95% confidence intervals. Only cells above the dotted lines were considered as expressing the mRNA and were used for the running window analysis.

We then wondered whether scaling could be detected as cells elongate and progress through a normal cell cycle. The fission yeast elongation phase is restricted to G2. Confounding effects arising from changes in genome content during DNA replication are therefore unlikely. Using cell length as a measure of size (**methods**), we analysed expression of the *rpb1* mRNA as a function of single cells’ sizes in *wt*, mutant and diploid cells and observed linear scaling within each genotype across a wide range of sizes (**Figure 1C and 1F**). Linear scaling during the cell cycle was apparent for all the constitutive genes analysed in this study (**Figure S1B-C**). This indicates that gene expression scaling is not an artefact of mutations in the *wee1* and *cdc25* cell-cycle regulators. Moreover, the mutant data demonstrate that gene expression scales over a dynamic range of sizes larger than that observed in *wt* cells. Together with earlier studies in cell populations this analysis establishes fission yeast as a powerful model system for studying scaling [3].

We next analysed the relation of scaling with ploidy using diploid cells. Diploids follow scaling parameters (slope, intercept) similar to smaller haploids (**Figure 1C, 1F**). Interestingly, for a given size, mRNA numbers in diploids and in *cdc25-22* haploids were similar (**Figure S1D-F**). Moreover, when analysing 3 diploid strains bearing heterozygous deletions of single genes we observed significant linear scaling of the corresponding mRNA but decreased concentrations compared to both diploid and *cdc25-22* cells. This suggests that scaling of single gene copies in fission yeast is coordinated with the cell genome content and that the machinery behind scaling may be limiting (**Figure S1D-F**).

At the level of single genes, is scaling a property of constitutive expression or do genomes of larger cells have a globally higher gene expression potential? To answer this question, we analysed three mRNA that are induced during specific phases of the cell cycle (**Figure S1G**). All three mRNA showed stronger induction levels in larger cells when comparing *wt* with *wee1-50* and *cdc25-22* mutants (**Figure 1D, 1G, S1I and S1J**). This indicates that larger cells have an increased gene expression potential in order to support scaling of constitutive mRNA and inducible transcripts, which were not expressed in smaller cells at the beginning of the cycle.

To investigate this further, we analysed the induction of two genes, which respond to acute changes in external conditions. The uracil regulated gene *urg1* responds to changes in uracil concentration and *sib1* to addition of 2,2-dipyridyl (DIP) [34, 35]. Induction of the *urg1* mRNA was heterogeneous and scaled with cell size across size mutants (**Figure 1E, 1H, and S1H**). The *sib1* mRNA showed a much more homogeneous response which also showed clear scaling within and across cell types (**Figure S1K**). This indicates that higher gene expression capacity in larger cells supports scaling in response to unexpected changes in external conditions. Taken together, these data indicate that scaling is universal and does not depend on the mode of gene regulation. Scaling does not result from a passive accumulation of mRNA during the cell cycle but from a change in the cell gene expression capacity that is coordinated with cell size.

### Coupling of mRNA decay rates with cell size is not a mechanism of scaling

Messenger RNA quantities are regulated at the level of transcription but also post-transcriptionally through modulation of degradation rates [36]. Three studies have reported that mRNA degradation rates are not regulated as a function of cell size in fission yeast, plant and mammalian cells [3,4,6]. To confirm these observations and extend them to single fission yeast cells, we analysed expression of 3 genes by smFISH after transcription inhibition with Thiolutin in *wt*, *wee1-50*, and *cdc25-22* cells. Thiolutin has been shown to inhibit transcription in *S. pombe* efficiently (**Figure S2A**) [11]. We observed mRNA half-lives of around 30-40min for the *rpb1* and *rpb2* mRNAs consistent with previous observations (**Figure S2B**) [37]. In *wee1-50* and *cdc25-22* mutants both mRNAs showed half-lives similar to *wt*, consistent with an absence of scaling of mRNA degradations rates. We also measured mRNA half-lives as a function of the cell cycle using cells binned by size. This analysis did not show consistent positive or negative coordination of degradation kinetics with cell size (**Figure S2B**, **left**). The absence of scaling of mRNA degradation rates was further confirmed using an orthogonal promoter switch-off approach for the *rbp1*, *rbp2* and *rtp1* genes (**Figure S2B**, **right**). Finally, as discussed in the next section mathematical modelling and inference do not support scaling of degradation rate. Overall, in agreement with previous studies, our analysis indicates that mRNA degradation is not a major mediator of scaling.

### Coupling of transcription rates to cell size and not burst frequency mediates scaling

We next explored the contribution of transcription rates to scaling using measurements of mRNA quantities and mathematical modelling. Using this combined approach allowed us to study dynamic transcription rates from static smFISH measurements of single cells. We developed agent-based models that incorporate the two-state model of gene expression inside growing and dividing cells, which are themselves described by phenomenological models of cell growth and size control (**Figure 2A, methods**). We used an Approximate Bayesian Computation (ABC) approach on cell size and smFISH measurements to infer mechanism of scaling. This inference approach determines the simplest model that captures the statistics of the experimental measurements and returns posteriors of the parameters and model probabilities for all models (**methods**). We used 2 classes of gene expression models that are the limiting cases of the two-state model shown in **Figure 2A**. The first class describes transcriptional bursts explicitly, while the second assumes only non-bursty transcription kinetics with a simple birth-death process that produces Poisson distributions (**Figure 2A**). Each class is in turn composed of two models the first assuming constant transcription rates during the cell cycle and the second assuming transcription rates that increase linearly (“scale”) with cell size (**methods**).

**Figure 2:**
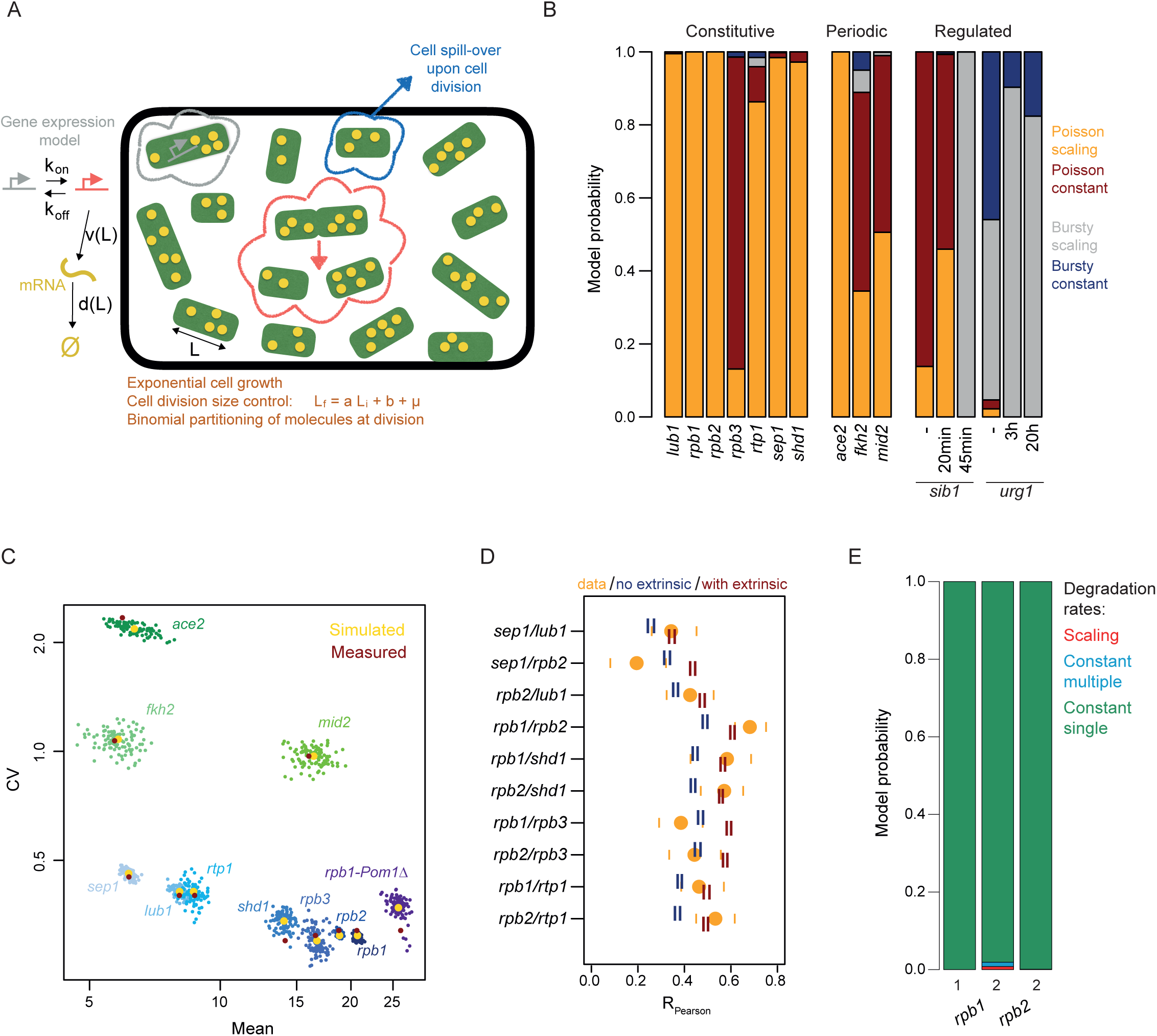
Coordination of transcription rates to cell size and not burst frequency nor mRNA decay rates mediates scaling. **A.** Cartoon of the modelling strategies used in this section (methods). Domino-like shapes represent cells/agents with mRNA pictured as yellow dots. Modelling of the cell cycle uses a noisy linear map framework and population size is kept constant by removing a random cell for every division event. The left part of the figures represents the two states model of gene expression. The transcription rate v and degradation rate d can be coupled to cell length (L). **B.** Bar chart showing the sum of particle weights for four models of transcription scaling after model selection analysis. Results for genes with three types of regulation are shown. Constitutive: constitutively expressed genes, Periodic: cell cycle regulated genes, Regulated: genes regulated by external stimuli. For regulated genes, times after addition of 2,2-dipyridyl (sib1) or uracil (urg1) are shown. The four types of model used in this analysis are marked on the right. Poisson: non-bursty transcription, Bursty: bursty transcription, Constant: constant transcription rates during the cell cycle, Scaling: transcription rates scaling with cell length (methods). **C.** Simulation of mRNA numbers during the cell-cycle using a model where transcription is scaling under a poisson regime. Coefficients of variation (CV) are plotted as a function of mean simulated mRNA numbers for each particle of each mRNA. Yellow circles denote the median of simulated data for all particles of a given mRNA. Red circles denote median for all experimental data used for parameter inference. **D.** Expression correlation between pairs of mRNA in single cells. smFISH experimental measurements are marked by orange dots with the 95% confidence interval shown with orange bars. 95% confidence intervals for correlations obtained from model simulations as in C are shown with blue bars. Red bars show the 95% confidence interval for simulations including 20% additional extrinsic noise (methods). **E.** Bar chart showing the sum of particle weights for three models of mRNA degradation after model selection analysis. Decay rate scaling: transcription constant and degradation rates scale with cell length, Constant single: *wt, wee1-50*, and *cdc25-22* cells have an identical constant degradation rate during the cell cycle with transcriptional scaling. Constant multiple: *wt, wee1-50*, and *cdc25-22* cells have distinct constant degradation rates during the cell cycle with transcriptional scaling.

We performed model selection between these four models on the smFISH data for 7 constitutively expressed genes in the *wt*. This analysis generated two clear conclusions.

First, models assuming bursty transcription were strongly penalised and supported by small model probabilities (**Figure 2B**, compare proportion of blue and red colours). Second, among the Poisson models, the one assuming transcription scaling with size was preferred for all but one mRNA (*rpb3* in **Figure 2B**, compare orange and dark red colours). From this we conclude that transcription rates of constitutively expressed genes in fission yeast are not bursty and scale with cell size.

We then asked whether transcription regulated by the cell cycle or by external cues followed the same paradigm. We found that, as constitutively expressed genes, cell-cycle regulated mRNAs do not follow a bursty transcription dynamics (**Figure 2B, middle**). This indicates that transcription, even when regulated in defined sections of the cell cycle falls into the Poisson regime. In terms of scaling, model selection is, as for the constitutive case, in overall support of scaling of transcription rate with cell size (**Figure 2B, middle**). Messenger RNAs that respond to external stimuli showed a different picture. Model selection favoured the bursty model with support for burst size scaling. This could point to bursty transcription of inducible genes, to the presence of strong extrinsic noise, or to a scenario where cells respond heterogeneously to the external signals.

In order to validate the model selection analysis, we generated simulations for all constitutive and cell-cycle regulated genes from **Figure 2B**, using models where transcription rates scale with cell size and follow a Poisson regime. This analysis reproduced quantitatively the mean and coefficient of variation (sd/mean, CV) of all experimental smFISH measurements, which illustrated the non-bursty nature of transcription for these genes further (**Figure 2C**). We then used the inferred parameters for the Poisson models to simulate gene expression of multiple genes in single cells and compared them to experimental data where 3 different mRNAs were measured in each cell (**methods**). This analysis suggests that most gene-to-gene correlations in expression are explained by scaling of transcription rates with cell size and may not require extensive additional regulation (**Figure 2D**). However, our data also suggest that for some gene pairs (e.g. Rpb1-Rpb2), correlations can be explained better by including some additional extrinsic variability.

We finally asked whether our modelling approach was also in support of a negligible role of mRNA degradation in scaling. We performed further model selection and simulation analyses on the transcription shut-off experiment from **Figure S2B** using models that consider different possible scalings of transcription and degradation rates (**methods**). This analysis shows that the model assuming transcriptional scaling with constant rates of mRNA degradation with cell size is overwhelmingly chosen (**Figure 2E**) and could capture experimental data quantitatively confirming our initial analysis (**Figure S2B**).

In summary, we show that, as in metazoans, scaling occurs through coordination of transcriptional rates with cell size in a rapidly growing unicellular organism. However, we find that transcription during the fission yeast rapid cell cycle is mainly Poissonian except during acute response to external changes where we detect signs of bursty expression. This is consistent with previous observation in the budding yeast *Saccharomyces cerevisae* [38]. Finally, we find that scaling of transcription rates explains most of the expression correlation of multiple mRNAs.

### Coordination of RNAPII initiation rates with cell size as the main mechanism of scaling

We then wondered which specific aspect of the transcription process was directly coordinated with cell size to mediate scaling. To investigate this and to confirm model predictions from **Figure 2**, we measured transcription rates experimentally. Single cells’ transcription rates can be estimated from smFISH images by measuring intensities of nuclear transcription sites [38]. As a model, we used probes directed against the 5’ region of the *rbp1* mRNA. This probe design provides strong sensitivity for detection of nascent transcripts (**Figure S3A, S3B**). Transcription site intensities and cell size were overall positively correlated when comparing *wt*, *wee1-50* and *cdc25-22* cells confirming modelling results and previous observation in a metazoan cell line (**Figure 3A, 3B, S3C**) [4]. We then investigated the impact of increased ploidy on transcription rates scaling. For this we compared nascent site intensities of 4 mRNA in haploid cells with those of diploid strains. We used probes against the 5’ end of *rpb1* as above and against three mRNA induced in specific phases of the cell cycle, since inducible expression increases the sensitivity of nascent site analysis. Intensities of individual transcription sites were similar between haploid and diploid cells (**Figure 3C**, green and yellow boxes). However, when considering the total intensity per cell, diploids showed increased rates and scaling was apparent again (**Figure 3C**, orange boxes). This indicates that the cell transcriptional capacity is limiting and distributed between the two gene copies of diploids. This was confirmed by analysis of the nascent intensity of the *rbp1* mRNA in a heterozygous deletion strain which showed no increased rates in the remaining copy (**Figure 3C**).

**Figure 3:**
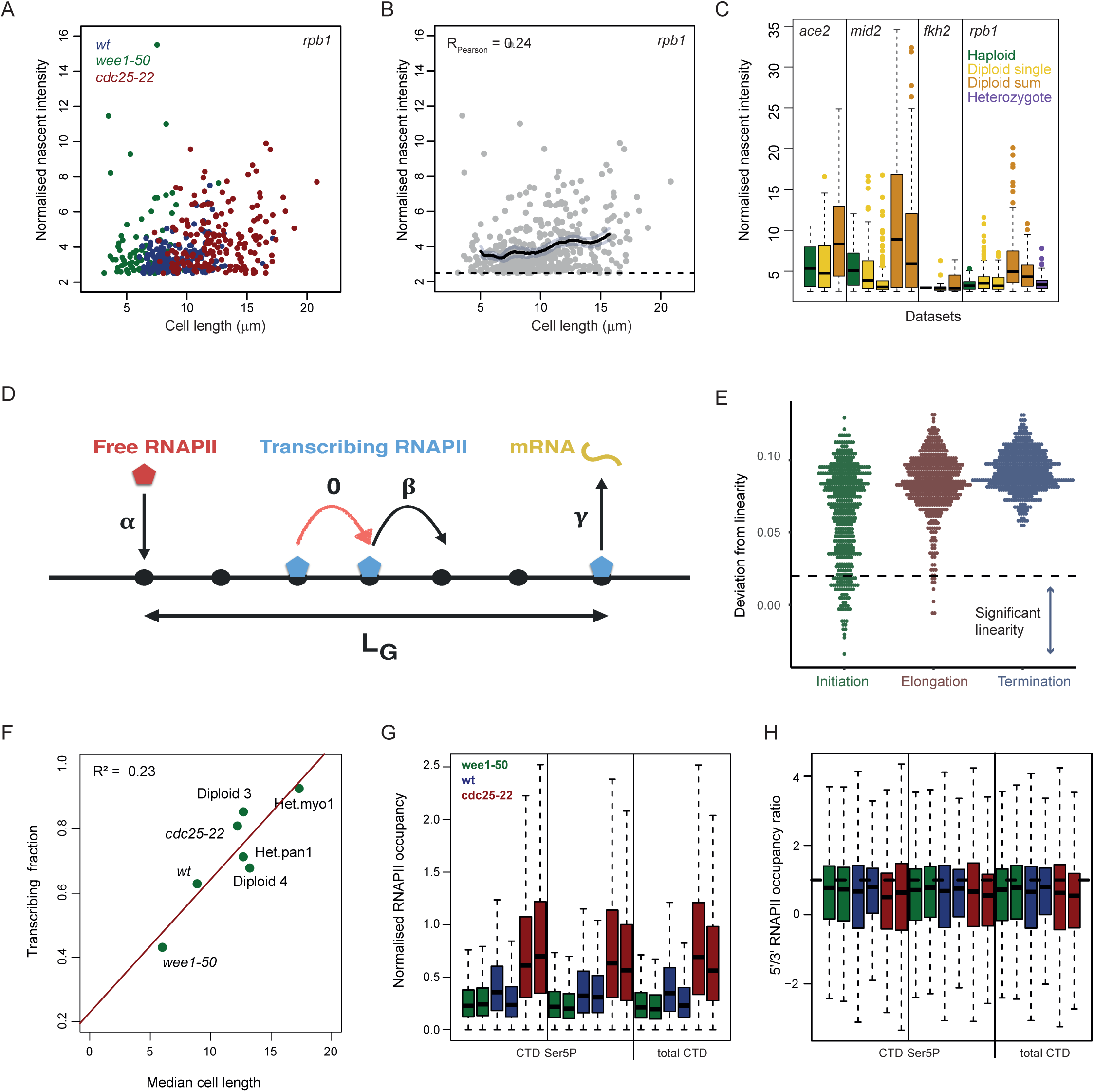
Coordination of RNAPII initiation rates with cell size as the main mechanism of scaling. **A.** *rpb1* normalised nascent sites intensities plotted against their respective cell length for *wee1-50* (green), *wt* (blue), *cdc25-22* (red). **B.** same as A. with all the cells in grey. The solid line shows median counts in running windows sampled from a count distribution identical to the experimental data. Shaded area represents 95% confidence intervals (methods). Cells with no nascent sites are excluded. Pearson correlation coefficient of nascent site intensity and length is shown. **C.** Boxplots of normalised nascent intensities for *rpb1* and the cell cycle regulated genes *ace2, mid2* and *fkh2*. Intensities are compared between haploid (green), diploid (yellow, orange), and *rpb1* heterozygote deletion cells (purple). For diploid cells the yellow boxplot shows intensities of single nascent sites, and orange boxplots show the total nascent intensities per cell. **D.** Cartoon of the TASEP modelling strategy of RNAPII transcription. A gene is represented by a lattice of length LG. RNAPII molecules enter the lattice at a rate α (Initiation), hop through the lattice at a rate β (Elongation) and exit the lattice at a rate γ (Termination, **methods**). **E.** Transcription linear scaling captured by the TASEP model where rates α (Initiation), β (Elongation) or γ (Termination) are linked to cell length. Simulations where run with each model with 500 random sets of TASEP parameters *α, β* and *γ* over times scales of 0.001 to 0.1 hours (**methods**). Deviation from linearity is shown for parameter samples in each of the 3 model variants. For each parameter set 1500 cells are simulated and two linear regressions are done on the mRNA numbers vs cell length data for the smaller and larger half of simulated cells. The deviation from linearity is estimated as the difference between the linear regression coefficients of small vs large cells. Values close to zero indicate a linear scaling and larger values indicate saturation of mRNA numbers at large cells. The dashed line shows the error in estimating the slope of the linear fit. Note that the initiation model shows the most robust linearity. **F.** Fraction of cells with a nascent site (“Transcribing fraction”) plotted as a function of mean cell length for the *rpb1* mRNA in a series of strains of different average length. Strains names are labelled next to the data points. Red line shows linear regression and R2 is shown in the top left corner. **G.** Chromatin immunoprecipitation analysis of RNAPII occupancy in wee1-50 (green), wt (blue), and cdc25-22 (red) cells. RNAP ChIP-seq data for antibodies against Serine 5 CTD phosphorylation (left, middle) and total Rbp1(right) are shown. RNAP occupancy values were normalised using DEseq2 and scaled using occupancy at histone genes (**methods**). **H.** RNAPII occupancy 5’/3’ ratio in wee1-50 (green), wt (blue), and cdc25-22 (red) cells. RNAP ChIP-seq data for antibodies against Serine 5 CTD phosphorylation (left, middle) and total Rbp1(right) are shown. RNAP occupancy values were normalised to input samples and each gene was divided into 100 bins using the deeptools package. Ratios between mean occupancy in bins 1-50 compared to bins 51-100 are shown (**methods**). Note that ratios are not changing with size and are consistently lower than 1.

To investigate further the mechanism behind scaling of transcription rates, we designed an orthogonal modelling approach where transcription is modelled as RNAPII particles hopping on a gene represented by a lattice (**Figure 3D**). This approach, which is based on a totally asymmetric simple exclusion process (TASEP), has been used successfully to study transcription and translation [6,39–42]. In our model, a gene is represented by a lattice of length *L_G_* and non-bursty transcription is modelled using three rates: i) the transcription initiation rate *α* is the rate at which RNAPII molecules enter the first site of the lattice (gene); ii) the elongation rate *β* is the rate at which RNAPII molecules hop one site forward on the lattice (gene); and iii) the termination rate *γ* is the rate at which the RNAPII molecule which sits on the last site of the gene leaves the lattice and produces a full length mRNA (**Figure 3D**). We incorporated this model in the agent-based framework from **Figure 2** assuming each rate could be linearly coupled to cell size.

By sampling the rates *α, β* and *γ* over physiological time scales estimated from previous studies (**Methods**) we found that coupling of initiation rates with size produced the most robust linear scaling (**Figure 3E**) and the strongest positive correlation of nascent intensities with size (**Figure S3D**). Although a model coupling transcription elongation rates with cell size could also generate linear scaling in some parameter regions (**Figure 3E**, **S3D**), these required elongation rates to be much slower than experimental measurements either in yeast or metazoans which report elongation rates of around 2kb/min (**Figure S3F**) [38,43– 47]. Importantly, linear scaling could only be observed in regimes with slow initiation rates relative to elongation and termination indicating that initiation is rate limiting (**Figure S3E**). Interestingly, this also suggests that fast, non-limiting initiation rates could be a mechanism by which some genes escape scaling (e.g. *rpb3*; **Figure 2B**). Finally, these results were confirmed by ABC inference analysis of nascent sites intensities data using the same TASEP model for 3 genes in different strains which showed clear preference for the initiation model (**Figure S3G**). This *in silico* analysis indicates that scaling of initiation rates with cell size is the likely mechanism of scaling.

The initiation model generated two important predictions. First, the model predicts that, even for non-bursty genes, cells in a population are not all actively transcribing at all times and the frequency of active transcription should increase with cell size. To test this prediction, we compared the fraction of cells with a nascent transcription site (“active cell fraction”) in *wee1-50*, *wt*, *cdc15-22* and diploid cells. As predicted by the model, a clear increase in the fraction of transcriptionally active cells with size could be observed (**Figure 3F**). Moreover, a strong positive correlation of the active cells fraction with cell length was apparent when calculated in sliding windows of increasing cell numbers during the normal cell cycle (**Figure S3I**). The second prediction of the initiation scaling model is a strong positive correlation between the number of transcribing RNAPII and cell size (**Figure S3D**). To test this, we analysed RNAPII occupancy across the genomes of *wt*, *wee1-50* and *cdc25-22* cells by chromatin immunoprecipitation followed by next generation sequencing (ChIP-seq). We used three antibodies against total RNAPII and Serine 5 phosphorylation of its carboxy terminal domain (CTD, **methods**). Using data normalised to occupancy at histone genes (**methods**) we could observe a significant increase in overall RNAPII occupancy with size consistent with previous observation and confirming model predictions (**Figure 3G**) [3]. Importantly, we did not find evidence for a redistribution of RNAPII from the 5’ to the 3’ of genes in larger cells which makes regulation of scaling at the level of RNAPII pause/release unlikely (**Figure 3H**, **S3J**). This profile was also reproduced quantitatively in our minimal TASEP model (**Figure 3D**, data not shown). In summary, our *in silico* and experimental data indicate that transcriptional scaling is mediated by an increase in RNAPII initiation rates coordinated with cell size. In addition, our modelling data together with the increase in RNAPII occupancy observed in larger cells suggest that RNAPII could be a limiting factor for transcription as cell size increases.

### RNAPII amounts on chromatin increases with cell size

If scaling of initiation rates is the mechanism behind scaling and RNAPII is limiting, the amount of RNAPII complexes on the genome of single cells should increase during the cell cycle. To test this hypothesis, we measured localisation of RNAPII in single cells by live-cell imaging. To do this, we tagged components of the RNAPII complex with green fluorescent protein (GFP) and imaged them during the cell cycle (**Figure 4A**, **S4A**). First, we observed that the cellular concentration of Rpb1 and Rpb9, two RNAPII subunits, remained constant during the fission yeast G2 growth phase (**Figure 4B**, **S4B**). This indicates that scaling of transcription initiation is not controlled by regulation of the cellular concentration RNAPII. Our image data show that a very large fraction of the tagged RNAPII subunits localises in the nucleus in the DAPI-positive area (**Figure 4A**). We therefore asked whether the amount of RNAPII on chromatin changes with cell size during the cell cycle. To assess this, we measured fluorescence intensities of the nuclear region occupied by RNAPII subunits (**Figure 4C**, **S4C**). Strikingly, the signal for both Rpb1 and Rpb9 increases steadily during fission yeast G2 phase where most cell size increase occurs (**Figure 4C**, **S4C**). Importantly this was not the case for another DNA bound protein the histone Hta2 (H2A beta), (**Figure. 4D-E**). This is consistent with the prediction of the initiation scaling model in the previous section and indicates that RNAPII quantities are likely to be limiting for transcription. In summary, these experiments indicate that increased initiation rates that mediate scaling are sustained by efficient import of RNAPII in the nucleus together with rapid recruitment onto chromatin.

**Figure 4:**
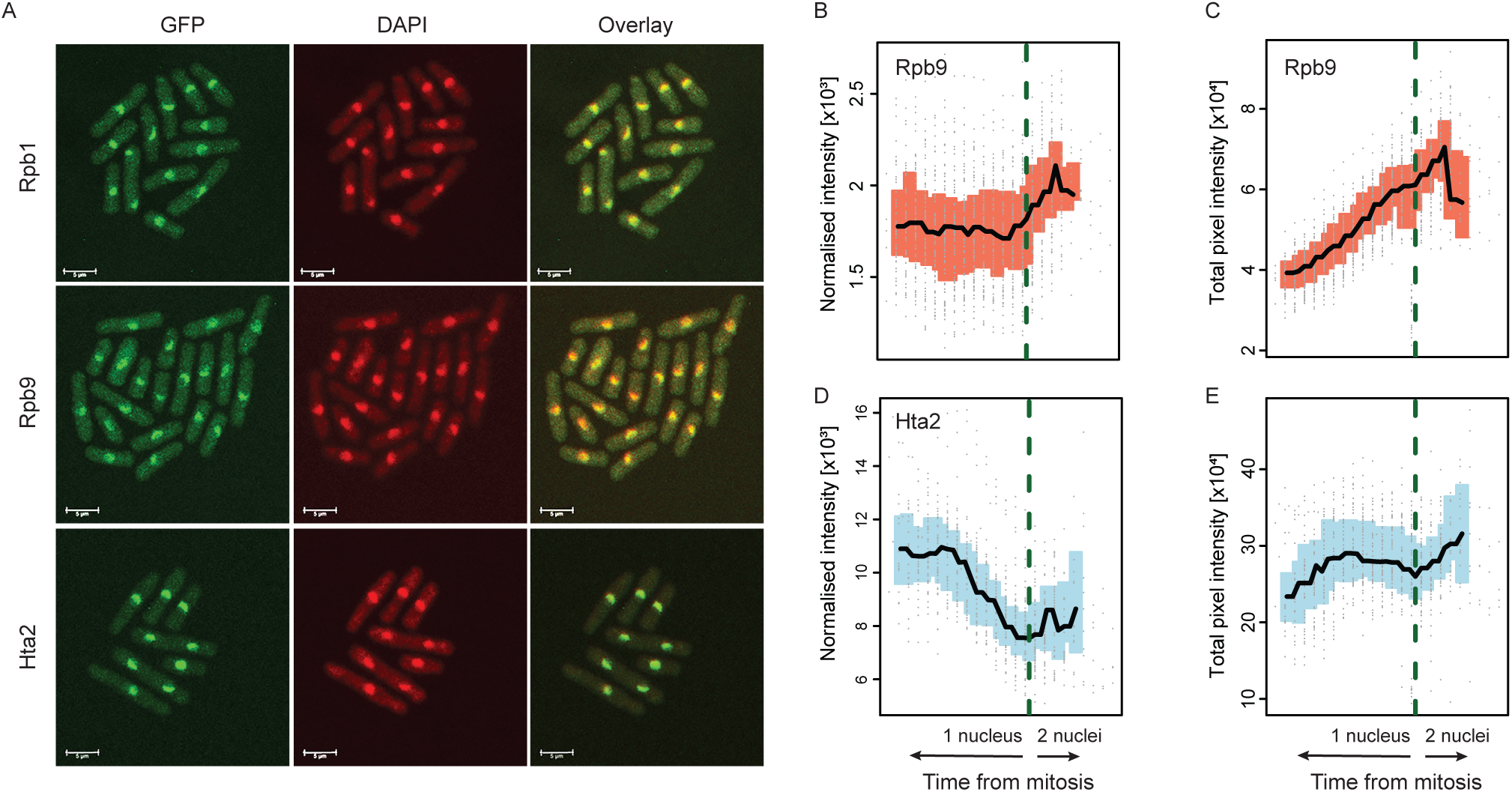
Nuclear RNA Polymerase II concentration increases with cell size. **A.** Confocal images of Rpb1, Rpb9, and Hta2 proteins tagged with GFP in wt fission yeast cells. DAPI straining for DNA and overlay of both channels are shown. The white scale bar represents 5µm. **B.** Widefield fluorescence data from live-cell imaging RNAPII subunit Rpb9. Normalised intensity/cell as a proxy for total cellular protein concentration is plotted along time relative to the mitotic phase of the cell cycle. **C.** Widefield fluorescence data from live-cell imaging RNAPII subunit Rpb9. Total pixel intensity in the area of strong fluorescence signal as a proxy for chromatin bound amounts is plotted along time relative to the mitotic phase of the cell cycle. **D.** As in B. but for the histone protein Hta2. **E.** As in C. but for the histone protein Hta2.

### The nucleus is the scaling unit

In fission yeast and other organisms, nuclear and cytoplasmic volume are intimately connected [48]. As scaling of initiation rates depends on RNAPII levels in the nucleus we wondered whether nuclear size rather than cell size itself could be the quantitative determinant of scaling.

To test this idea, we analysed nascent site intensities of *cdc11-119* mutant cells cultivated at non-permissive temperature. Under these conditions, cells elongate, undergo mitosis and nuclear division but do not divide (**Figure 5A**) [49]. Strikingly, scaling of nascent site intensities of individual nuclei with cell size was not apparent in this system (RPearson = 0.06, **Figure 5B**, **right**). Scaling was only restored when all nascent sites present in a cell were added together (R_Pearson_ = 0.46, **Figure 5B**, **left**). This mirrors data from multinucleated cells showing that while the overall nuclear volume scales with cell volume, the volume of individual nuclei is proportional to their surrounding cytoplasmic volume [48]. Consistent with this, the ratios of nascent site intensities between nuclei of *cdc11-119* cells were weakly correlated with their immediate cytoplasmic volumes (**Figure S5A**). This indicates that the cell “scaling unit” could be the nucleus and not the whole cell.

**Figure 5:**
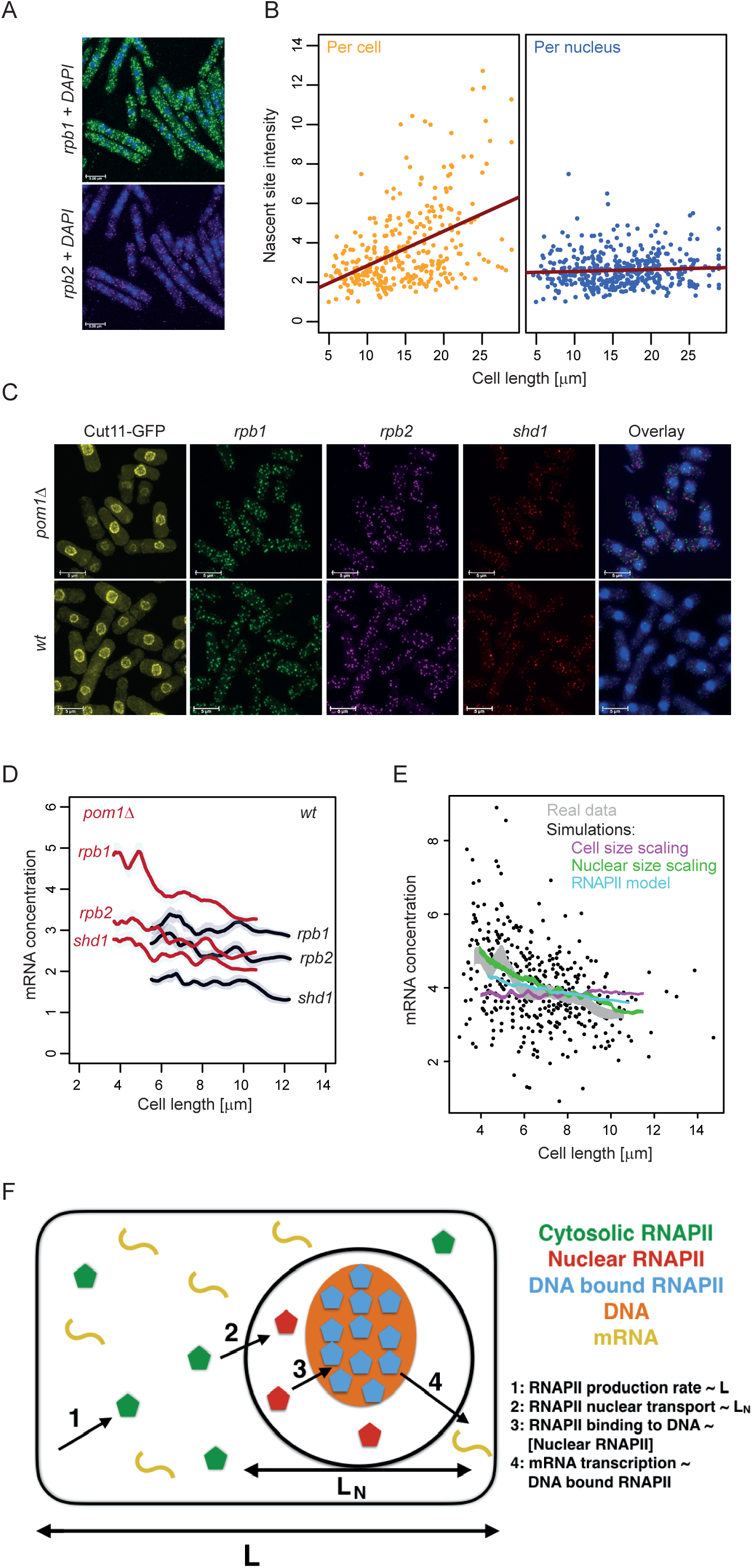
The nucleus is the scaling unit. **A**. Representative smFISH images for the rpb1 and rpb2 mRNA in cdc11-119 grown at non-permissive temperature. The white scale bar represents 5µm. **B**. rpb1 normalised nascent sites intensities per nucleus (blue) or per cell (orange) plotted as a function of each cell respective size. Red lines are regression lines from linear models. The white scale bar represents 5µm. **C**. Representative smFISH images for the *rpb1*, *rpb2*, and *shd1* mRNA in *wt* and *pom1* deletion mutants carrying the Cut11-GFP nuclear marker (left column). Overlay is shown on the right. The white scale bar represents 5µm. **D**. mRNA concentration (mRNA/cell length) for the *rpb1*, *rpb2*, and *shd1* mRNA in *wt* (black) and *pom1* deletion mutants (red). The solid line shows median counts in running windows sampled from a count distribution identical to the experimental data. Shaded area represents 95% confidence intervals. **E**. *rpb1* concentrations measured by smFISH are plotted as a function of cell length. Running average for the experimental data (grey) or derived from model predictions assuming transcription rates scaling with cell length (magenta), nuclear area (green), or predicted from a mechanistic model of RNAPII prediction are shown (light blue). **F**. Cartoon representation of a mechanistic model of RNAPII transcription (see Methods for details and parameters).

We next analysed scaling in conditions where the correlation between cell and nuclear size is compromised. Pom1 is a regulator of cell polarity and division which when deleted leads to increased size variability at cell birth due to cell partitioning errors [50]. We analysed expression of three mRNA by smFISH in *pom1Δ* mutant cells expressing a marker of the nuclear envelope to allow measurement of nuclear size (**Figure 5C**). As expected cell and nuclear size show smaller correlation in the *pom1Δ* mutants compared to *wt* cells (**Figure S5B**). Scaling in this system was comparable to *wt* cells with the exception that *pom1Δ* mutants showed a higher y-axis intercept when mRNA numbers are plotted as a function of cell size (**Figure S5C**). This indicates that mRNA concentrations in *pom1Δ* cells are higher in smaller cells after birth and negatively correlated with cell size (**Figure 5D**). This deviation from perfect concentration homeostasis is also observed but to a lower degree in *wt* cells (**Figure 5D**) as reported for mammalian cells [4]. Moreover, mRNA numbers divided by nuclear volume show a negative correlation with nuclear volume (**Figure S5D**). The modelling approach described in **Figure 2** which is based on transcription scaling with cell size and binomial partitioning of mRNAs based on daughter cell sizes failed to capture this behaviour (**Figure 5E**, **magenta line**). However, a modified model coupling transcription rates to nuclear size instead produced a good fit to the data (**Figure 5E**, **S5F green line**). Together, this analysis suggests that beside the nucleus being the scaling unit, nuclear size could be an important determinant of scaling.

### A mechanistic model of scaling

We used the results from this study to develop a mechanistic model of scaling centred around RNAPII mediated transcription (see also [19, 20]). (**Figure 5F**, **methods**). In this model the rates of RNAPII complex synthesis and maturation scale with cell size. This is consistent with our smFISH and live-cell imaging data showing that RNAPII subunits have a constant cellular concentration during the cell cycle (**Figure 1 and 4**). RNAPII is then transported to the nucleus with a rapid rate that is depleting cytosolic RNAPII (as observed in live-cell imaging data; **Figure 4)**. Once in the nucleus, RNAPII binds to DNA with a constant high affinity and transcription rates are proportional to the numbers of DNA-RNAPII complexes present on each gene at any given time consistent with scaling of initiation rates (**Figure 3**). Finally, RNAPII levels is set to be limiting in line with the initiation scaling model (**Figure 3**), the behaviour of diploid and heterozygous mutants (**Figure 3**) [10] and with the fact that the cell synthetic capacity is titrated against the number of genes in heterokaryons [4]. Moreover, this assumption fits the observation that many RNAPII subunits are limiting for growth in fission yeast [51–53]. The high affinity and the limiting amount of RNAPII ensures that the majority of RNAPII molecules are bound to DNA and increase with cell size consistent with our imaging data (**Figure 4**) and with biochemical evidence from mammalian cells [4]. Finally, DNA replication occurs close to cell division and each daughter cell inherits about half of the DNA-bound RNAPII mostly independent of its size as observed in our live cell imaging experiment (**Figure 4C**, **S4C**). This simple model is able to capture the different features of our data. First, it retrieved the scaling of DNA-bound RNAPII with cell size while keeping the overall cellular concentration constant as observed in **Figure 4** (**Figure S5E**). Second, it explained the scaling of mRNA numbers with cell size including the higher mRNA concentration observed in smaller cells (**Figure 5E**). As this mechanistic model relies on both cell size mediated production of RNAPII and nuclear mediated partitioning of RNAPII it captures better the negative correlation between mRNA numbers over nuclear area and cell size than a purely nuclear size mediated transcriptional scaling (**Figure S5E**, compare blue and green line). Overall, this simple mechanistic model of RNAPII transcription by integrating the experimental findings from this study, explains the origin of transcriptional scaling and its subtler features and identifies the competition between genes for the limited pool of transcriptional machinery as central to the phenomenon of transcriptional scaling.

## CONCLUSIONS

We performed an extensive experimental and modelling study of gene expression scaling with cell size in single fission yeast cells. We found that scaling is a pervasive feature of gene expression that impinges on constitutive and regulated expression. We then showed that scaling relies on an increase in RNAPII initiation rates with cell size and a concentration independent recruitment of a limiting RNAPII on the genome. Finally, we propose that the nucleus is the scaling unit and that nuclear size may participate in setting scaling levels.

Our work supports a simple and robust model for the scaling of gene expression with cell size in which the competition between promoters for a limited pool of RNAPII determines their relative strength. Because RNAPII maintains a constant concentration as cell size increases (as proteins do in general), the number of RNAPII complexes increase linearly with cell size. cell size increase will not affect the relative strength of promoters, but will cause their absolute rate of transcriptional initiation to scale linearly with cell size, exactly as required to produce the observed gene expression scaling.

Our model assumes that the general and specific transcription factors that regulate relative promoter strength are in excess even in small cells, and thus are not affected by cell size. To gain a deeper mechanistic understanding of scaling, it will be important to determine whether this assumption is met for most regulators or if some escape the rule. It could also inform about mechanisms through which some mRNA escape regulation by scaling as our modelling results suggest that non-limiting initiation rates can produce this behaviour [10–12]. Another interesting question will be to determine the role of chromatin remodellers in facilitating transcription initiation in larger cells. It is possible that a more permissive chromatin environment in large cells synergises with increased RNAPII local concentrations to support higher transcription rates of inducible genes.

Recent evidence suggests that, in mammals, burst size is regulated at the level of the proximal promoter sequence, while distal enhancers are involved in setting burst frequency [54]. Moreover, burst initiation and RNAPII pause/release but not RNAPII recruitment have been shown to be regulated in response to biological perturbations [55]. Our model of scaling through initiation of RNAPII transcription fits well with these data as this process is independent of both gene activation and control of burst frequency. It is also consistent with the observation that promoters maintain their relative expression levels in changing growth conditions [56].

Our observation that scaling is regulated at the level of single nuclei in multinucleated cells and may be linked to nuclear size is interesting. RNA synthesis levels have been connected to nuclear size in other systems such as multinucleated muscle cells [57], or for the HTLV-1 mRNA [58]. It could reflect a higher availability of RNA polymerases around larger nuclei as these tend to be surrounded by larger cytoplasmic volume (**Figure S5A**) [48, 57]. Interestingly, the nucleus was also found to be an independent transcriptional unit in mature osteoclasts [59] and in multinucleated fungi where nuclei retain local control of cell cycle periodic transcription [60, 61]. This suggests that more complex feedback and molecular mechanisms may also be at play.

An important result from our study, which is not directly related to scaling, is the absence of bursty transcription for most fission yeast genes tested. This is in line with previous observations in budding yeast and plants [29,38,62]. Transcriptional bursts result in high gene expression noise and are associated with Fano factors (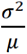 of mRNA numbers) greater than 1, whereas Poissonian birth-death processes have a Fano factor = 1. Our finding that most transcription followed a non-bursty regime relied on our modelling taking cell size and the cell cycle into account explicitly. Without doing so, all genes in this study have been thought to have bursty expression as Fano factors calculated on the raw counts were well above 1 (not shown). This reiterates the importance of studying gene expression considering potential confounding effects of morphological features such as cell size and the cell cycle [23,24,29,63,64].

Finally, in addition to progressing our understanding of the mechanisms behind scaling, this study provides a large quantitative dataset of gene expression and cell size measurement in over 20000 cells in various conditions. This will support future modelling efforts aimed at understanding regulation of gene expression.

## ACKNOWLEDGEMENTS

We would like to thank Nick Rhind for stimulating discussion and his help with formulating our model of scaling. We are grateful to Kurt Schmoller, Philipp Thomas, Marc Sturrock and Nick Rhind for critical reading of the manuscript. We thank Snezhana Oliferenko and Jürg Bähler for sharing strains. This research was supported by the UK Medical Research Council, a Leverhulme Research Project Grant (RPG-2014-408). It used the computing resources of the UK Medical Bioinformatics partnership (UK MED-BIO), which is supported by the UK Medical Research Council (grant MR/L01632X/1) and the Imperial College High Performance Computing Service. ChIP-seq data have been deposited in ArrayExpress, accession number XXXXXX.

## METHODS

### Strains and culture conditions

The strains and their genotypes that were used in this study are listed in **Table S1**. Genetic crossing confirmed by polymerase chain reaction was used for strains generated unless otherwise specified. Strains were revived from glycerol frozen stocks on solid yeast extract agar (YE agar), or YE agar supplemented with 25 mg l^-1^ adenine, L-histidine, L-leucine, uracil, L-lysine, and L-arginine (Sigma), and with or without appropriate antibiotics for selection. YE agar plates were incubated for approximately 48 h at 32°C in a static incubator until visible large colonies could be observed. Single colonies were transferred into liquid yeast extract medium (YE), in YE supplemented as above (YES), Edinburgh minimal medium (EMM), or EMM supplemented as above (EMMS), unless otherwise indicated in figure legends, and incubated at 170 rpm in a shaking incubator. Temperature sensitive strains were grown at 32°C and shifted to 36.5°C for the time indicated in figure legends. For the induction of *sib1* expression, the strains were grown at 25°C to an optical density at 600 nm (OD600) of ≈0.4 and treated with 2,2-dipyridyl (DIP; ACROS) at a final concentration of 250 µM for the time indicated in figure legends, or left untreated. For measuring mRNA decay rates, cells were grown in YE at 25°C to OD600 ≈0.4; cells were treated with thiolutin (AXXORA) for the time indicated at a final concentration of 15 µg/ml, or left untreated. For *urg1* induction, cells were grown in EMM supplemented with or without 0.25 mg/ml uracil for the time indicated. For transcription inhibition, log phase cultures (OD600∼0.5) were treated with thiolutin (15 ug/ml) and same volume of DMSO (used for dissolve thiolutin) was added to thiolutin untreated culture. Samples were taken at 0, 25, 35, 45 mins and processed as for smFISH. For live-cell experiments, cells were grown in EMMS in syphonstats – chemostat-like devices (http://klavinslab.org/hardware.html) which maintain the turbidity of liquid cultures by diluting with fresh medium appropriately – and maintained at OD600 0.4 at 32°C by frequent dilution [65].

### RNA single molecule fluorescence in situ hybridisation (smFISH)

All smFISH datasets are described in **Table S2**. All the mRNA counts, nascent site intensities and cell size measurements are available in **Table S3**. smFISH samples were prepared according to a method modified from published protocols [66, 67]. Briefly, cells were fixed in 4% formaldehyde and the cell wall partially digested using zymolyase. Cells were permeabilised in 70% ethanol, pre-blocked with bovine serum albumin and salmon sperm DNA, and incubated overnight with custom Stellaris oligonucleotide sets (Biosearch Technologies) labelled with CAL Fluor Red 610, Quasar 670, or Quasar 570 (probe sequences are listed in **Table S4**). Cells were mounted in ProLong™ Gold antifade mountant with DAPI (Molecular Probes), and imaged on a Leica TCS Sp8 confocal microscope, using a 63x oil objective (NA 1.40). Optical z sections were acquired (0.3 µm step size) for each scan to cover the entire depth of cells. Cell boundaries were outlined manually and single mRNA molecules were identified and counted using the FISH-quant MATLAB package [68]. Cell area, length, and width were quantified using custom ImageJ macros. The technical error in FISH-quant detection was estimated at 6–7% by quantifying *rpb1* mRNA foci with two sets of probes labelled with different dyes. The nascent mRNA foci were identified and quantified using intensity information from the above-mentioned FISH quantification where an intensity 2.5 to 3-fold above the modal intensity within the same cell was chosen as a threshold for nascent mRNA. The quality of the identification of nascent sites was validated manually by visualising high intensity foci in the nucleus, with an accuracy of over 90% in all three strains (wild-type, *cdc25-22*, and *wee1-50*).

### ChIP-seq

Chromatin immunoprecipitation (ChIP) assays was carried out essentially according to published methods [69]. In brief, cells were grown in YES to an OD600 of ∼ 0.8 and fixed with formaldehyde solution (1% final) and then quenched with glycine. After washing twice with cold PBS (phosphate-buffed saline), cells were re-suspended in lysis buffer containing proteases inhibitors and disrupted vigorously with acid-washed glass beads 8-11 times for 20 sec in a FastPrep instrument. Samples were then sonicated in Bioruptor (at High setting and 6 times 5 mins with 30 secs ON/30 secs OFF). Chromatin were immunoprecipitated with antibody against Rpb1 (ab817) or Rpb1 CTD-ser 5 (ab5408 or sigma 04-1572), which were coupled to Dynabead protein-G and protein-A and Dynabead sheep anti-mouse or rat IgG (Invitrogene). DNA was purified from immunoprecipitated samples using MinElute Qiagen kit. Quantification of the DNA was done using QuDye dsDNA HS assay kit and quality was verified using Bioanalyzer.

For sequencing DNA from immunoprecipitated samples, the libraries were made using the NEBNext ChIP-Seq Library Prep Master Mix Set for Illumina (E6240S) with the indexes provided in NEBBext Multiplex Oligos for Illumina (Index Primers Set1,2 and 3). Negative control DNA are those from the same chromatin extracts without going through immunoprecipitation steps. Pools of libraries were sequenced on an Illumina HiSeq 2500 instrument at the MRC LMS genomics facility. Paired-end reads (100 nt) were generated from two pools of 12 or 18 samples per sequencing lane. Data were processed using RTA 1.18.64, with default filter and quality settings. The reads were de-multiplexed with bcl2fastq-1.8.4 (CASAVA, allowing 0 mismatches).

### ChIP-seq analysis

A description of ChIP-seq libraries can be found in **Table S5**. Sequencing reads were aligned to the fission yeast genome as available in PomBase in July 2019 using BWA [70, 71]. For figure 3G, RNAPII occupancy counts were extracted for each transcript using HTseq [72] and the fission yeast annotation available in PomBase in July 2019 [71]. Data were normalised using DESeq2 and scaled using the mean counts of fission yeast histone genes as a scaling factor to allow comparison of global RNAPII occupancy between size mutants [73]. Amounts of histone proteins are thought to scale with DNA content rather than cell volume and are commonly used as normalisation factors for absolute proteomics measurements [74]. Moreover, average synthesis rates of histones were found to remain constant across a wide range of sizes in budding yeast (Kurt Schmoller personal communication). For figures 3H and S3I, RNAPII immunoprecipitation data were normalised with their respective input and average gene analysis was performed using the deeptools analysis suite [75].

### Live-cell microscopy and analysis

Strains of interest and wild-type ySBM2 were grown from single colonies in 5 ml YES before they were transferred into syphonstats and maintained at OD_600_ 0.4 overnight in EMMS. Prior to microscopy, ySBM2 cells were mixed at a 1:10 ratio with each strain of interest and diluted to a final OD600 of 0.3 in fresh EMMS. Cells were loaded directly into a CellASIC^®^ ONIX Y04C-02 microfluidic plate (EMD Millipore) according to the manufacturer instructions. Fresh EMMS was continually perfused through the growth chamber with a constant pressure of 6.9 kPa (approximately 3 µl/h). Cells were imaged on an Olympus IX70 inverted widefield fluorescence microscope with an environmental chamber maintained at 32°C, with a high precision motorised XYZ stage (ASI), controlled with µManager version 1.4.22 [76]. Cells were continually imaged at a 10 minute interval with a 40× objective (NA 0.95, UPlanSApo; Olympus) with brightfield (30 ms exposure), GFP (250 ms exposure, emission filter Semrock 514/30 nm), and dsRed (500 ms exposure, emission filter Semrock 617/73 nm) channels captured by a Hamamatsu Orca Flash 4.0 V2 sCMOS camera, with illumination provided by a Lumencor Spectra X LED light source set to 20% power. For each of the four growth chambers within the microfluidic plate, three positions were defined, each of which was focussed using the software autofocus using the brightfield channel; since this feature was generally inaccurate, a 3 µm Z-stack was used around the autofocus position to ensure that at least one Z-position was in focus for each position.

Initial analysis was performed with Fiji – a derivative of ImageJ – with time-lapses assembled if necessary, and in-focus slices for each time-point selected using a custom macro (written by Stephen Rothery, Facility for Imaging by Light Microscopy, Imperial College London), resulting in a 4-dimensional OME-TIFF file for each field-of-view (XYCT). Each file was subsequently analysed using a series of custom Python scripts utilising the scikit-image, SciPy, NumPy, and pandas packages extensively among others. The scripts permit the semi-automated definition of cell boundaries (segmentation) within the brightfield channel, followed by the quantification of fluorescence within cell boundaries, identification of nuclei, further quantification within nuclear boundaries, and assignment of cells into lineages. Cell segmentation was effected using a custom ‘balloon-filling’ algorithm, in which a connected series of nodes is ‘inflated’ from its centre inducing an outwards force on all nodes; this outward force is counteracted by an ‘image force’, which applies an opposing force inversely proportional to the intensity of pixels neighbouring the node, this has the effect of preventing nodes from expanding through areas of low light intensity – which generally surrounds the cell boundary; finally each node is also affected by its direct neighbouring nodes which pull each node sideways according to their position, ensuring smooth contours. Together, after multiple iterations, nodes migrate from a central location until the forces equilibrate at areas of low light intensity (generally the edge of the cell). This procedure can be performed in an automated manner in which iterations cease when the area contained within the nodes does not significantly change, or in a supervised manner, in which further iterations are prompted manually via a keyboard command, with progress displayed via a graphical interface. Initial centres of cells are defined either by manually clicking, or by the centre of the cell in the previous frame. Nuclei are defined as areas within the cell boundary which have red fluorescence pixel intensity values greater than 1.1 standard deviations above the mean intensity within the whole cell boundary. Time from mitosis is defined as the number of hours from the point at which the number of detected nuclei increases from one to two. Fluorescence intensity is adjusted for uneven illumination according to a series of empty fields imaged using the same settings. Background and autofluorescence is determined from wild-type cells cultured within the same field-of-view, with the mean level of fluorescence within these cells subtracted from measurements. Fluorescence is normalised by cell or nuclear area by dividing total fluorescence of all pixels within their respective boundary by the area of that boundary. Scripts are available upon request.

For live cell imaging in Figure S4, cells were imaged in ibidi microfluidic channel slides (μslide VI 0.4, #80601) instead of a CellASIC ONIX microfluidic plate. 30 μL of cells sampled from syphonstat cultures maintained at OD600=0.5 for at least 15 generations were loaded into channels of a pre-warmed slide. 40 μL of pre-warmed EMMS was then added to each reservoir of channels, followed by 10 μL of mineral oil (Sigma, M5904) to prevent evaporation during imaging. Live-cell microscopy was performed as described above.

Image analysis was semi-automated and performed using custom interactive MATLAB scripts available upon request. Tracks corresponding to individual cell cycles are recorded by user clicking. Only cell cycles for which cells remained in in focus for at least the 2 hours preceding mitosis were collected. Local background subtraction was automatically applied to fluorescence images. Cell segmentation was performed automatically based on thresholding of the brighfield images. Nuclear segmentation was performed automatically based on thresholding of the uch2-mCherry fluorescence images.

### Cell size measurements

We extracted both cell area and cell length measurements form the smFISH images as proxies for cell volume. We observed that both measurements support robustly data characteristics such as scaling of mRNA numbers and positive intercepts. We used cell length as a proxy for cell volume throughout the manuscript as it proved to be a simpler and more consistent measure. Importantly, as fission yeast has a cylindrical shape, its length is directly proportional to its volume. For the nucleus, we acquired area measurements only. As the nucleus is spherical, area and volume are not proportional. We have therefore derived volume estimates from area measurements assuming a perfect sphere using:

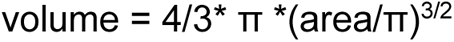

### Mathematical modelling

We use agent-based simulations of stochastic gene expression coupled to cell size in growing and dividing cells (**Figure 2A**) [24]. We assume cells grow exponentially with a constant growth rate from birth to division that is sampled from a truncated Gaussian distribution with mean *m_g_* and standard deviation *σ_g_*. A cell that is born with birth length *Li*grows until it reaches the division length *L_f_*. We use a phenomenological model of cell size control that relates the final size to initial size through a noisy linear map, which captures experimentally observed variability and correlations in cell size [77–79].

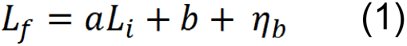

where *a* and *b* are size control parameters (*a* = 0 denotes a sizer mechanism and *a* = 1 an adder mechanism) and *η_b_* is a truncated Gaussian with mean zero and standard deviation *σ_b_*. The dividing cell of length *L_f_* produces two daughter cells with birth sizes 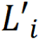 and 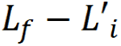, where 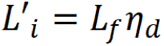 and *η_d_* is another truncated Gaussian with mean 0.5 and standard deviation *σ_d_*. The biomolecules such as mRNA molecules (except for DNA) are binomially partitioned in the daughter cells with a probability proportional to the daughter cell size

As shown in **Figure 2A**, we simulate a fixed number of cells, so upon cell division one of the existing cells (including one of the newly born daughter cells) is chosen randomly and taken out of the simulation. This ensures we are simulating a constant number of cells in time and can produce snap-shot data with the correct cell age and size distribution as observed in the experimental data. This has been used in the simulations used in the ABC inference (**Figure 2**). For the modelling results shown in **Figure 3 and 5**, where the results are conditioned on cell size, we have used a simpler scheme [24], where upon cell division only one of the daughter cells is followed modelling a single lineage (similar to the data generated in a mother machine) [80]. The simulations are performed using a simple algorithm that uses discretised time steps to simulate exactly the Gillespie method [81] with time-dependent parameters [82]. The simulation code for inference is written in the Julia programming language and the rest of simulations are performed in R. The codes are available upon request.

Our main gene expression model is the so-called telegraph model or the two-state model [83] where the gene can be in an ‘off’ or an ‘on’ state and transcription can only occur when the gene is on (**Figure 2A**). If the gene is always on then this model reduces to a simple non-bursty birth-death process with parameters transcription rate *v* and mRNA decay rate *d*. Here the mRNA counts per cell have a Poisson distribution (in the absence of cell cycle effects). In the limit where the duration of the promotor on-state is shorter than the mRNA life time we have the bursty limit characterised by 3 parameters of average burst frequency *k_on_*, average burst size *v/k_off_* and mRNA decay rate *d*. In this model, birth events are simulated as geometrically distributed increases in mRNA numbers [84]. For **Figure 2B** model selection, we used four variants of this model including the Poisson and bursty limits with or without transcriptional dependence to cell size. Transcriptional scaling is modelled as linear dependence of transcription rate or burst size to cell size (*v*(*L*) = *v*_0_*L*, where, *v*_0_ is a constant and *L*,is the cell length). The models for the cell cycle regulated genes, assume that there is a point in the cell cycle, where gene expression increases from a basal level to an active level. ABC model selection in **Figure 2E** is performed on the data from the transcription shut-off experiments of the 3 strains of *wt*, *wee1-50*, and *cdc25-22.* The 3 models included are all the Poisson limit with different scaling assumptions for transcription and decay rates. Model one assumes transcriptional scaling with a single constant decay rate across the 3 strains, the second model assumes also transcriptional scaling but with 3 different constant decay rates within each strain. And the third model assumes constant transcription and decay rate that is proportional to inverse cell size across the 3 strains. The priors used in the model selection, are wide over a physiological range. The model selection results were not too senstivie to the choice of the priors.

In **Figure 3D** a more detailed model of transcription is illustrated, which is based on the totally asymmetric simple exclusion process (TASEP) [6,39–41]. Here, the promoter is assumed to be always active, i.e. we are modelling a non-bursty gene. The gene is modelled as a lattice of size *L_G_*. Transcription starts by initiation through binding of a RNAPII molecule to the first site on the gene with rate *α* if the site is empty. Elongation is modelled as hopping of RNAPII forward with rate *β* if the next site is available. Termination is modelled with RNAPII leaving the last site on the gene with rate *γ* which gives rise to a fully transcribed mRNA. In our model, we have ignored pausing, backtracking and incomplete termination. In **Figure 3E**, we compare 3 variants of this model, where size scaling is through linear coupling of initiation, elongation or termination rate to cell size. We chose *L_G_* = 20 for computational efficiency and as it is larger than the typically observed number of RNAPIIs on the genes, which is related to nascent site intensity (**Figure 3A**). In each model, we randomly picked the initiation time scale *τ_1_*, elongation time scale through the whole gene *τ_E_*, and termination time scale *τ_T_* between 0.001-0.1 hours that produces average mRNA numbers of between 20-30 for a moderately expressed gene. The lower limit on the time scale is significantly shorter than the mRNA life time and the upper limit represents very slow steps to achieve moderate transcription, given the mRNA life time, and is also slower than the time-scales reported in the literature [cite]. The rates are inversely proportional to the time-scales as *α* = 1/*τ_1_*, *β* = *L_G_*/*τ_E_* and γ = 1/*τ_T_* Note that as *β* is the rate of hopping per site, it is also proportional to the number of sites on the gene *L_G_*. We also performed an ABC model selection using an implementation of our TASEP model in Julia on several datasets (**Figure S3G**). We used the same prior as discussed above.

The nuclear scaling model and the RNAPII model in **Figure 5E** rely on both cell size and nuclear size dynamics. It is known that nuclear size scales closely with cell size [48]. There has not been much modelling of nuclear size control in the literature. We introduce a phenomenological and passive model of cell and nuclear size control, extending the noisy linear map of cell size control (**Equation 1**). We assume cellular exponential growth, cell size control and division as before. We assume nuclear size also grows exponentially and follows its own noisy linear map:

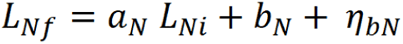

Cell division time is determined when cells reach their final size (*L_f_*) For simplicity, we assume mitosis is taking place at cell division and the size of the newly divided daughter cells and their nucleus is determined by 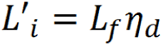 and 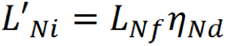, where *η_d_* and *η_Nd_* are truncated correlated Gaussian noise with mean equal 0.5, standard deviations *σ_d_* and *σ_d_* and correlation coefficient of *ρ_d_*. We choose *ρ_d = 0.5_* based on analysis of our time-lapse imaging data (**Figure 4**) and the rest of the parameters of our dual noisy linear map model of cell and nuclear control were fitted on the static pom1 mutant size data using the ABC inference.

Given, our dual noisy linear map model discussed above, in the nuclear scaling model (**Figure 5E**), we assume transcription rate *v* to be linearly dependent on the nuclear size *L_N_* but mRNAs are partitioned upon division based on the size of the daughter cells (not nuclear size of the daughter cells). In this model a small daughter cell is likely to inherit a nucleus of average size, with transcription rates higher than expected from cell size, resulting in an increase in mRNA concentration for small cells.

The RNAPII model (**Figure 5F**) provides a mechanistic RNAPII based model of transcriptional scaling. In this hybrid deterministic and stochastic model, transcription, translation, complex formation and maturation of RNAPII molecules are modelled as simple cell size dependent production steps. The RNAPII is then transported to the nucleus by a nuclear size dependent rate and it binds to DNA with high affinity with a rate that is dependent on the concentration of DNA in the nucleus (inversely proportional to nuclear size). In this model transcription rate of a gene is assumed to be proportional to the amount of DNA-bound RNAPII.

In this hybrid deterministic and stochastic model, RNAPII dynamics are modelled deterministically by a series of ODEs inside growing and dividing cells, while transcription of mRNA is modelled stochastically. Upon cell division, we assume mRNA are partitioned binomially according to the size of the daughter cells. The free cytosolic and nuclear RNAPII are portioned binomially according to the size of the daughter cells and their nucleus. The scaling in this model comes about from sequestration of RNAPII on the DNA. The model is very robust to different model parameters and assumptions as long as the level of free cytosolic and nuclear RNAPII is much smaller than the DNA bound RNAPII. The qualitative model results for Rpb1 shown in **Figure 5E**, are obtained by using the parameters of the dual noisy linear map model discussed above for the pom1 mutant, tuning RNAPII parameters to obtain about 10% free RNAPII and linear scaling of DNA bound RNAPII, as well as setting the transcription rate to match expression levels of Rpb1. The model without any further tuning recovers deviation from concentration homeostasis observed at small cell sizes, which is observed for the different genes in the WT and Pom1 mutant strains.

### ABC inference and model selection

In this study we have used Approximate Bayesian Computation (ABC) for inference. When the likelihood function is intractable, we require a tool for carrying out inference without it. ABC is precisely such a tool. The algorithm originated in the 1980s and 90s (see e.g. [85]). For review of more recent developments, see [15]. ABC aims to carry out Bayesian inference without having to evaluate the likelihood function. Given data *D* and model with parameter set *θ,* this is done by approximating the posterior distribution:

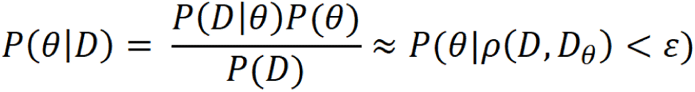

Where *D_θ_* is a set of data generated from the model with parameters *θ*, sampled from the prior *P*(*θ*), *ρ*(*D, D_θ_*) is a distance measure that is defined on the set of such datasets (or their summary statistics) and *ε* is a tolerance, representing the degree of approximation we are willing to accept. The simplest ABC algorithm, that is based on sampling *θ* repeatedly and rejecting the ones that produce data with larger distance than our tolerance (which is called ABC rejection sampling [86]), is too inefficient. Much work has been carried out over the past decade in this area, leading to a variety of different implementations with much more favourable scaling of computation time with the dimensionality of the parameter space [86]. For the purpose of this project we will use a Sequential Monte Carlo implementation, based on the implementations of Toni et. al [87] (ABC-SMC) and Lenormand et.al [88] (APMC). In the ABC-SMC, one fixes the size of the posterior sample, *N*, and a finite sequence of decreasing tolerances, {*ε_t_*}, a priori. The primary differences between APMC and ABC-SMC are firstly that the sequence of epsilons is not determined a priori; it is dynamically determined from the previous iteration’s distribution of errors until a stopping criterion (*pacc_min_*) is fulfilled and secondly that simulations from earlier iterations are not discarded. ABC lends itself very naturally to model selection [86, 87]. In essence, all we have to do is to extend our priors to one extra dimension, representing different models. Formally, we require a joint prior distribution over models and parameters, *P*(*M, θ*). We have combined the model selection aspects of ABC-SMC implementation and adaptive aspect of APMC to obtain our APMC with model selection algorithm.

In order to apply our APMC algorithm, we need to choose an appropriate distance *ρ*(*D, D_θ_*). As the problem at hand is stochastic in nature, we have chosen to use sum of square differences of summary statistics of the data and simulated data in the distance measure:

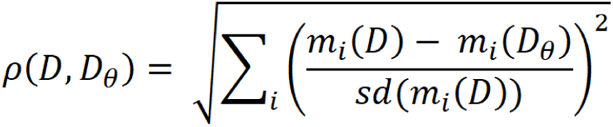

where, we used central sample moments and cross moments *m_i_*(*D*) of our data up to order 3. With our data sample sizes, moments beyond the first three are usually too noisy to be useful. Also, each term in the distance measure are weighted by the bootstrap estimates of standard deviation of the central moments. This rescales the terms in the sum appropriately and downweighs the noisier moments, helping to prevent overfitting of the data.

---

**Figure S1 related to Figure 1.**
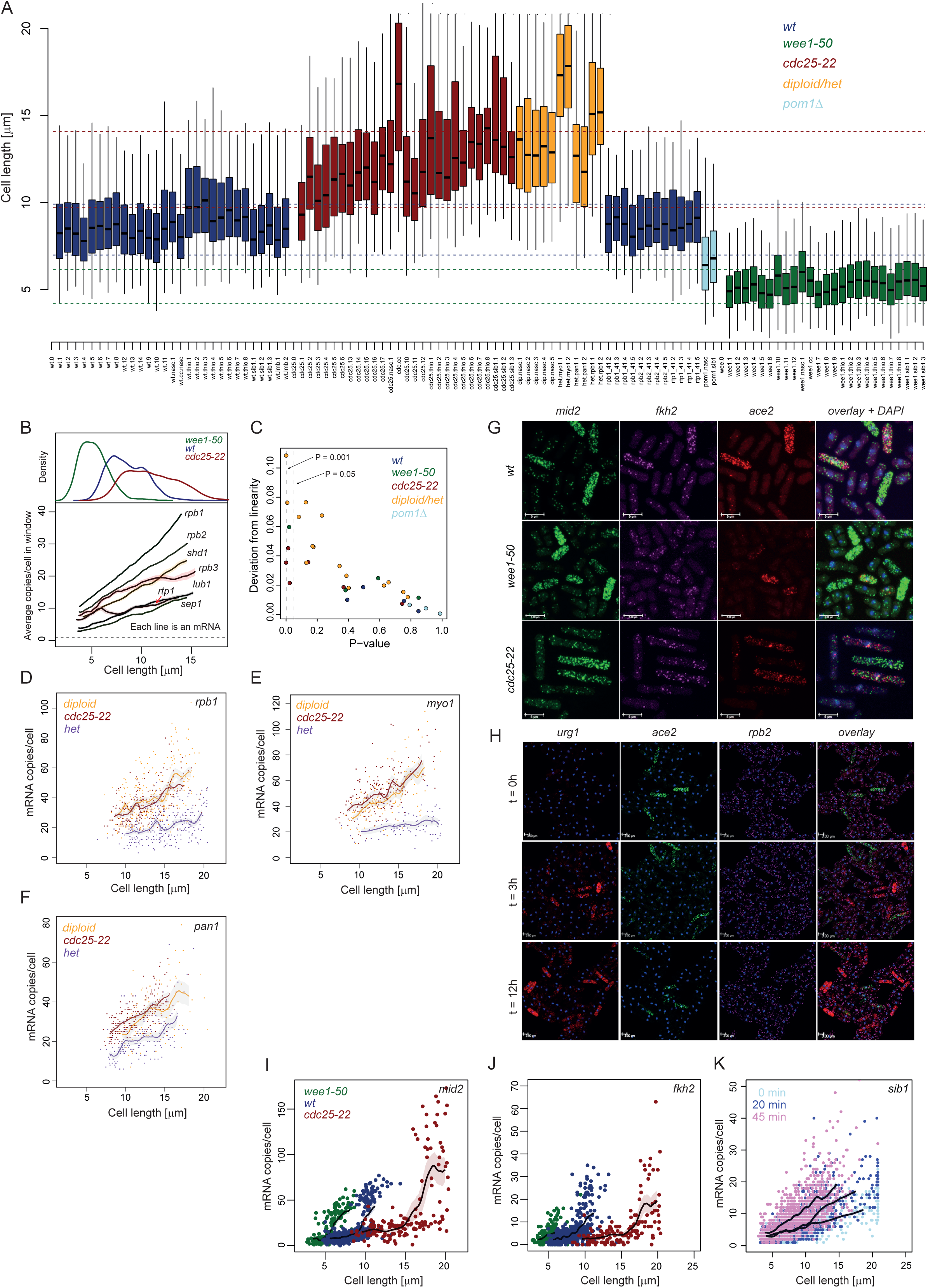
**A.** Cell length distribution for the 111 datasets described in this study. Colours represent different genotypes. Datasets are described in **Table S2**. Coloured lines show the interquartile range for all the *wee1-50* (green), *wt* (blue), *cdc25-22* (red), diploid and heterozygote deletions (orange) and *pom1*Δ (light blue) cells. **B.** Top: Pooled count distribution of 7 constitutively expressed genes in *wee1-50* (green), *wt* (blue), and *cdc15-22* (red) cells. Bottom: Median counts of 7 constitutively expressed genes as a function of cell length in running windows sampled from a count distribution identical to the experimental data. Shaded area represents 95% confidence intervals (**methods**). Data from *wee1-50*, *wt*, *cdc15-22* cells are pooled and gene names are indicated next to their respective data line. **C.** Deviation from linearity is plotted as a function of linearity P-values (see Figure 3E, **methods**). Low P-values mean significant deviation from linearity. Colours are as in panel A. P-values equal 0.05 and 0.001 are shown with dashed lines and annotated. **D.** *rpb1* mRNA copies/cell plotted against cell length for diploid (orange), *cdc25-22* (red) and *rpb1* heterozygote diploid (purple). The solid line shows median counts in running windows sampled from a count distribution identical to the experimental data. Shaded area represents 95% confidence intervals (**methods**). **E.** *myo1* mRNA copies/cell plotted against cell length for diploid (orange), *cdc25-22* (red) and *myo1* heterozygote diploid (purple). The solid line shows median counts in running windows sampled from a count distribution identical to the experimental data. Shaded area represents 95% confidence intervals (**methods**). **F.** *pan1* mRNA copies/cell plotted against cell length for diploid (orange), *cdc25-22* (red) and *pan1* heterozygote diploid (purple). The solid line shows median counts in running windows sampled from a count distribution identical to the experimental data. Shaded area represents 95% confidence intervals (**methods**). **G.** Representative images of an smFISH experiment for cell cycle regulated genes. The *mid2*, *fkh2*, and *ace2* mRNA are labelled in *wee1-50*, *cdc25-22* and *wt* cells. Overlay of the three channels and DAPI staining of DNA is shown on the last column. The white scale bar represents 5µm. **H.** Representative images of an smFISH experiment for a gene regulated by external conditions (*urg1*). The *urg1*, *ace2*, and *rpb1* mRNA are labelled in *wt* cells before, 3 hours and 12 hours after addition of uracil to the culture. Overlay of the three channels and DAPI staining of DNA is shown on the last column. The white scale bar represents 5µm. **I.** *mid2* mRNA copies/cell plotted against cell length for *wee1-50* (green), *wt* (blue), cdc25-22 (red). The solid line shows median counts in running windows sampled from a count distribution identical to the experimental data. Shaded area represents 95% confidence intervals (**methods**). Only cells above the dotted lines were considered as expressing the mRNA and were used for the running window analysis. **J.** *fkh2* mRNA copies/cell plotted against cell length for *wee1-50* (green), *wt* (blue), cdc25-22 (red). The solid line shows median counts in running windows sampled from a count distribution identical to the experimental data. Shaded area represents 95% confidence intervals (**methods**). Only cells above the dotted lines were considered as expressing the mRNA and were used for the running window analysis. **K.** *sib1* mRNA copies/cell plotted against cell length for wt cells before (light blue), 20min after (blue) or 45min (pink) after addition of 2,2-dipyridyl (DIP). The solid line shows median counts in running windows sampled from a counts distribution identical to the experimental data. Shaded area represents 95% confidence intervals (**methods**).

**Figure S2 related to Figure 2.**
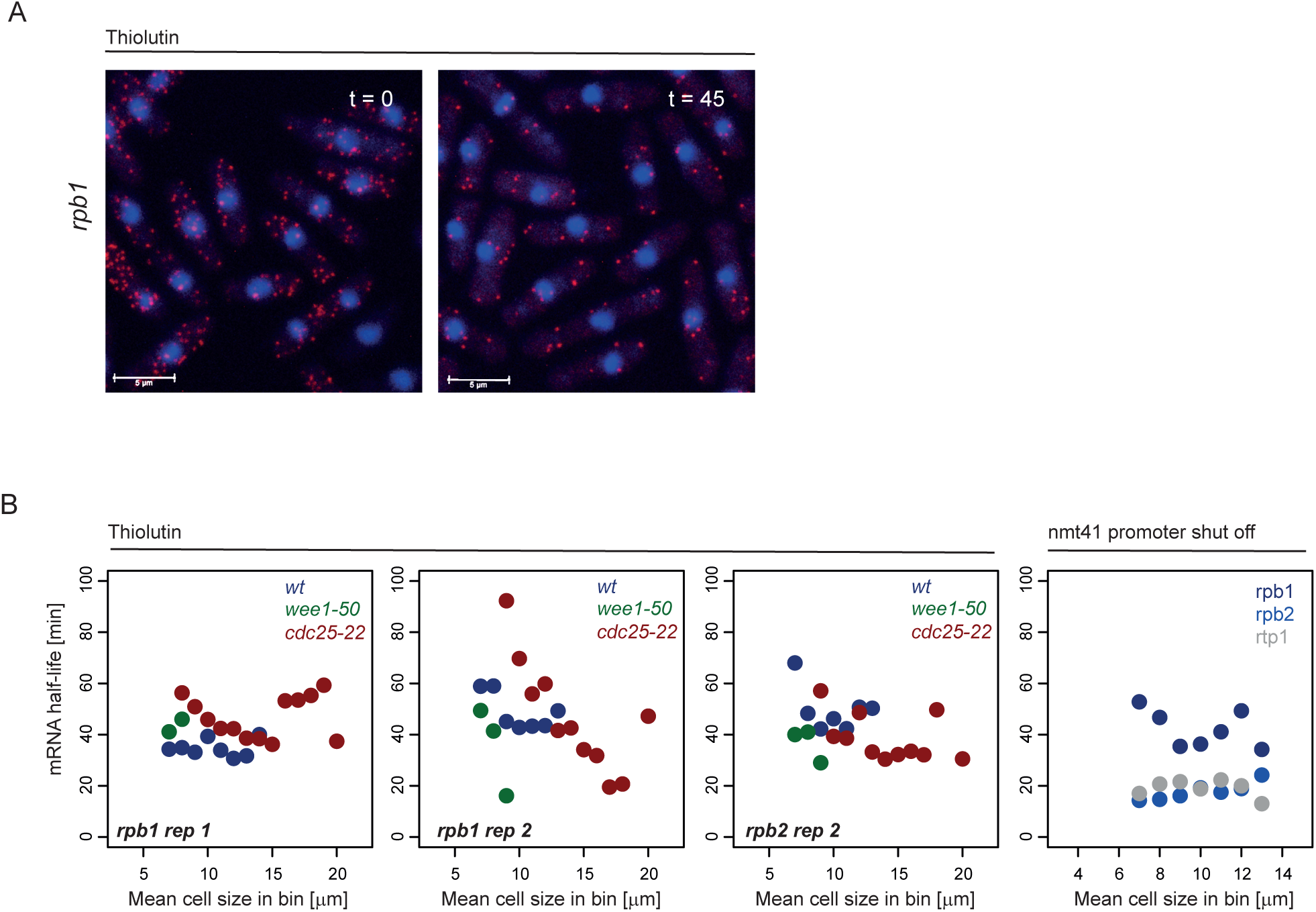
**A.** Representative images of smFISH experiments for the *rpb1* mRNA in cells before (t = 0) and after a 45min (t = 45) treatment with 15 µg/ml of Thiolutin. Note the number of mRNA dots decreases with time and that the transcription foci disappear after treatment. The white scale bar represents 5µm. **B. Left:** mRNA half-lives were measured after Thiolutin treatment in bins of increasing cell length for the *rpb1* (2 repeats) and *rpb2* mRNA in *wee1-50*, *wt*, and *cdc25-22* cells (left). **Right:** mRNA half-lives were measured after nmt41 promoter switch-off in bins of increasing cell length for the *rpb1*, *rpb2* and *rtp1* mRNA in *wt* cells.

**Figure S3 related to figure 3.**
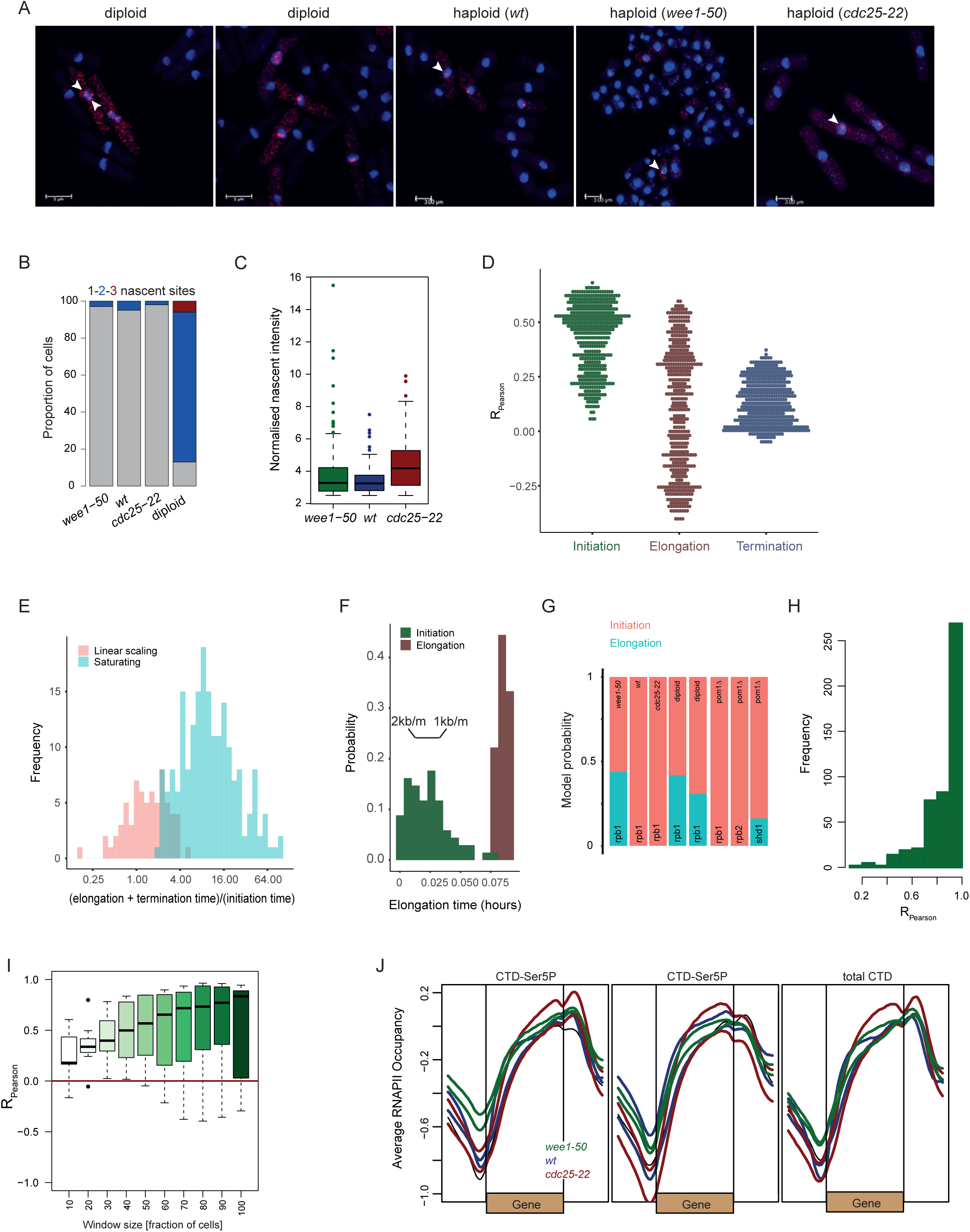
**A.** Representative images of smFISH experiments for the *rpb1* mRNA in haploid and diploid cells. Nascent transcription sites are indicated with white arrows. The white scale bar represents 5µm. **B.** Fraction of cells with 1, 2, or 3 nascent sites (as in A) in haploid and diploid cells. Note the diploid cells have mostly 2 sites validating our approach specificity and sensitivity to detect nascent transcription sites. **C.** Boxplot of nascent sites intensities for the *rpb1* gene in *wee1-50* (green), *wt* (blue) and *cdc25-22* (red) (related to Figure 3A). **D.** Pearson correlation between mRNA numbers and cell length for the simulation data from Figure 3E. **E.** Histogram of ratio of initiation time scale (1/*α*) and the sum of elongation time scale and termination time scale 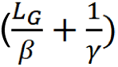 for parameter sets resulting in linear scaling and parameter sets resulting in saturating transcription scaling indicating that initiation rate should be limiting for linear transcriptional scaling. **F.** Elongation time of a 2kb gene obtained from the simulations from Figure 3E for the models with Initiation rates (α, green) and elongation rates (β, red) coupled with size. Only simulation that achieved linearity are shown. Elongation time from published experimental measurements of elongation rates (1-2kb/min) are shown. Note that the elongation scaling simulations require elongation rates much slower than experimentally measured in order to achieve linearity. **G.** ABC model selection results between the initiation scaling and elongation scaling TASEP models for 3 genes across several strains. **H.** Distribution of the correlations of frequencies of actively transcribing cells with cell length for different parameter sets from the Figure 3E simulations. Note the strong positive correlations as observed in experimental data (Figure 3F). **I.** Boxplot of correlations between normalised intensities of the rpb1 mRNA and cell length for all cell genotypes in sliding windows or containing different number of cells. Each box represents a window of a given width. Sliding windows are shifted with increment of 1 cell along all cells ordered by length. Note that correlations are mainly positive. **J.** Average gene analysis for the RNAPII ChIP-seq data from Figure 3H. Data were normalised to input samples and split in 200 bins (50: upstream, 100: gene, 50: downstream) using the deeptools package.

**Figure S4 related to figure 4.**
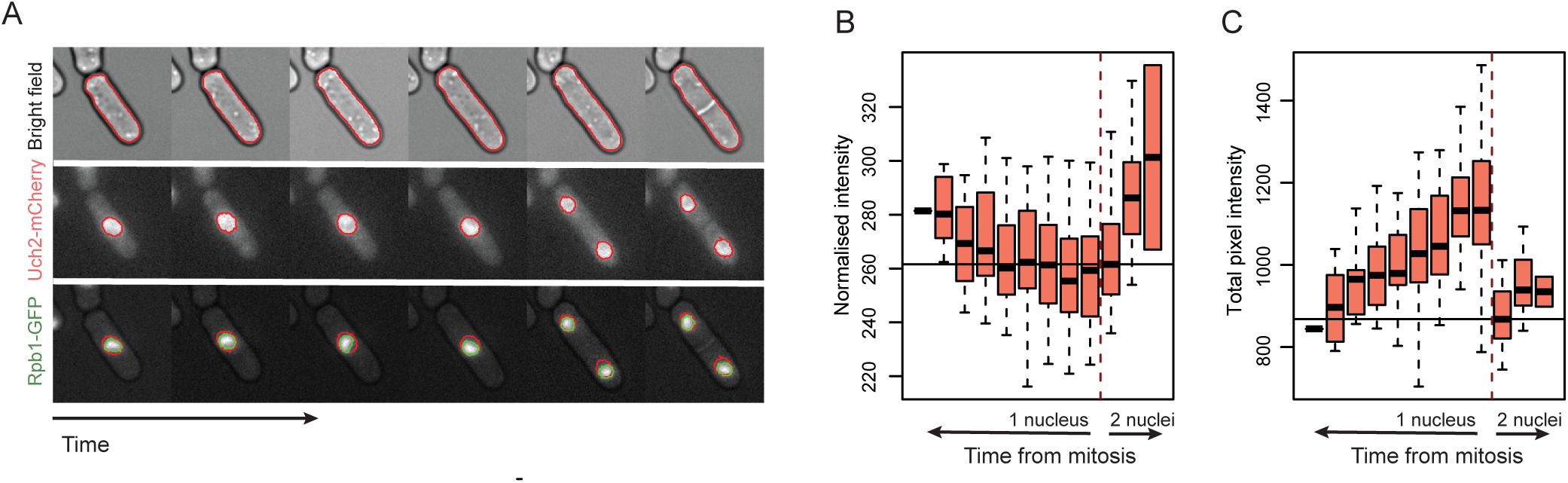
**A.** Representative image of a single cell followed for a full cell-cycle using ibidi slides. Brightfield (top), the nuclear marker Uch2 (middle) and the RNAPII subunit Rpb1 bottom. **B.** Rpb1 normalised intensity/cell as a proxy for total cellular protein concentration is plotted along time relative to the mitotic phase of the cell cycle. **C.** Rpb9 total pixel intensity in the area of strong fluorescence signal (green mask in A) as a proxy for chromatin bound amounts is plotted along time relative to the mitotic phase of the cell cycle.

**Figure S5 related to figure 5.**
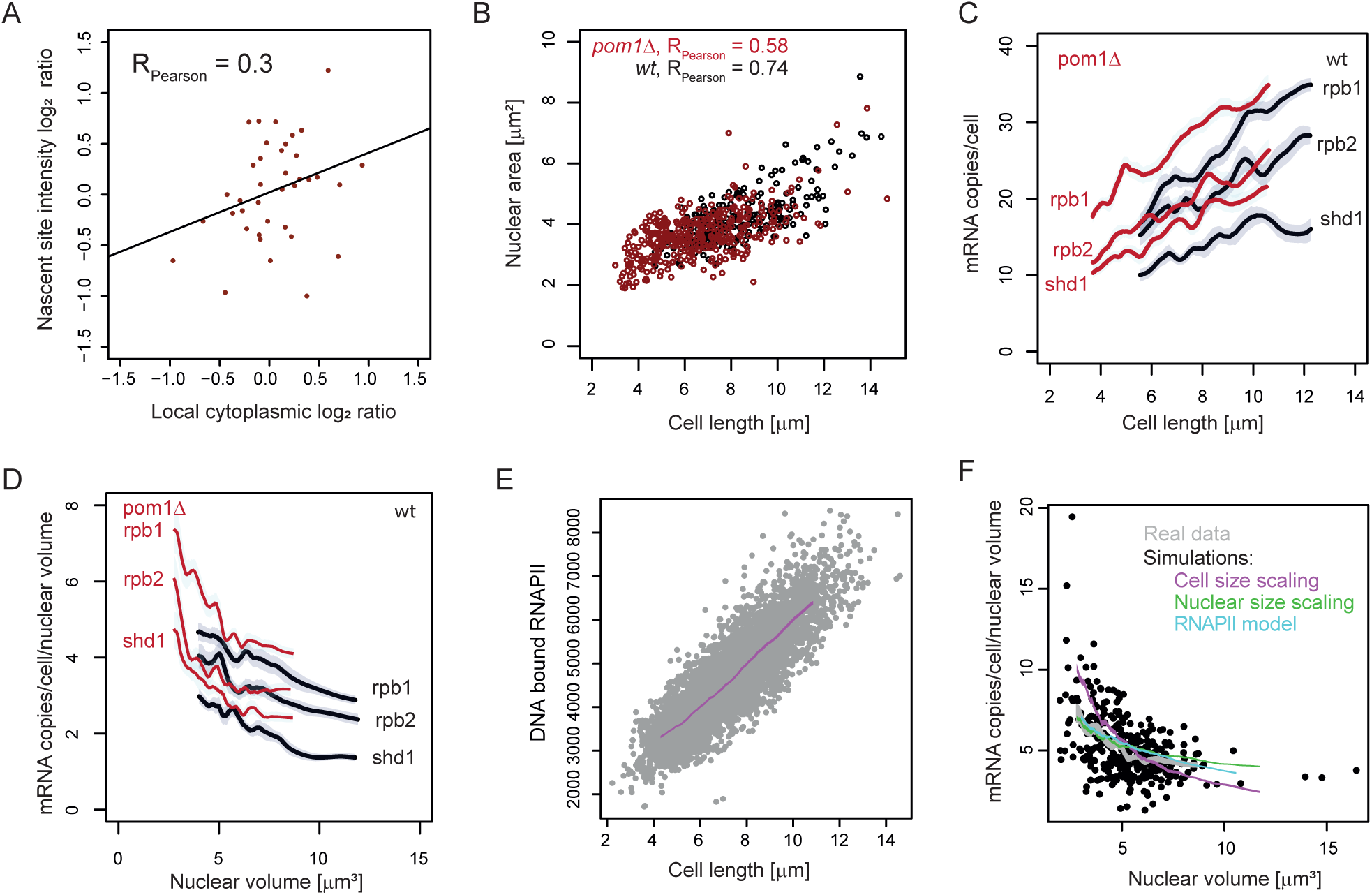
**A.** Ratios of nascent site intensities of bi-nucleated cdc11-119 cells plotted as a function of the ratio of local cytoplasmic volumes (**methods**). The black line is the regression line from a linear model. **B.** Correlation between cell length and nuclear area in pom1Δ (red) and wt (black) cells. Pearson corretions are shown. **C.** As in Figure 5D for mRNA numbers. **D.** As in Figure 5D for mRNA numbers/nuclear volume plotted as a function of nuclear volume. **E.** Simulation data from the model described in Figure 5F. DNA bound RNAPII is plotted as a function of cell length. Magenta lines show a running average. **F.** As in Figure 5E for mRNA numbers divided by nuclear area.

